# Functional Characterization of *fer-ts*, a Temperature-Sensitive FERONIA Mutant Allele That Alters Root Hair Growth

**DOI:** 10.1101/2020.05.27.119891

**Authors:** Daewon Kim, Jiyuan Yang, Fangwei Gu, Sung Jin Park, Jonathon Combs, Alexander Adams, Heather B. Mayes, Jeong Dong Bahk, Erik Nielsen

## Abstract

In plants, root hairs undergo a highly-polarized form of cell expansion called tip-growth, in which cell wall deposition is restricted to the root hair apex. In order to identify essential cellular components that might have been missed in earlier genetic screens we identified conditional temperature sensitive (ts) root hair mutants by EMS mutagenesis. Here we describe one of these mutants, *fer-ts* (*<u>fer</u>onia-<u>t</u>emperature <u>s</u>ensitive*). Mutant *fer-ts* seedlings grew normally at normal temperatures (20°C), but failed to form root hairs at elevated temperatures (30°C). Map based-cloning and whole genome sequencing revealed that *fer-ts* resulted from a G41S substitution in the extracellular domain of *FERONIA* (*FER*). A functional fluorescent fusion of FER containing the *fer-ts* mutation localized to plasma membranes, but was subject to enhanced protein turnover at elevated temperatures. While tip-growth was rapidly inhibited by addition of RALF1 peptides in both wild-type and *fer-ts* mutants at normal temperatures, root elongation of *fer-ts* seedlings was resistant to added RALF1 peptide at elevated temperatures. Additionally, at elevated temperatures *fer-ts* seedlings displayed altered ROS accumulation upon auxin treatment and phenocopied constitutive *fer* mutant responses to a variety of plant hormone treatments. Molecular modeling and sequence comparison with other CrRLK1L receptor family members revealed that the mutated glycine in *fer-ts* is highly conserved, but is not located in the recently characterized RALF23 and LORELI-LIKE-GLYCOPROTEIN (LLG2) binding domains, perhaps suggesting that *fer-ts* phenotypes may not be directly due to loss of binding to RALF1 peptides.

## INTRODUCTION

In higher plants, root hairs are cellular protuberances resulting from the polarized outgrowth of specialized root epidermal cells, known as trichoblasts (Gilroy and Jones, 2000). Root hair development can be divided into three phases: cell specification, initiation of bulge formation, and polarized tip growth (Cho and Cosgrove, 2002). Polarized tip growth results from highly localized exocytosis of Golgi-derived vesicles containing newly synthesized cell wall material at a restricted area of the root hair tip, with a tip-focused cytoplasmic calcium ion (Ca^2+^) gradient ensuring correct targeting of this polarized membrane trafficking (Hepler et al., 2001; Smith et al., 2005; Cole and Fowler, 2006). This Ca^2+^ gradient is established by localized generation of reactive oxygen species (ROS) by *ROOT HAIR DEFECTIVE2* (*RHD2*) which encodes an NADPH oxidase in *A. thaliana* (Foreman et al., 2003). RHO OF PLANTS (ROP) small GTPases mediated signal transduction is involved in specifying the root hair initiation site, and future root hair elongation by stimulating *RHD2* activity at the growing root hair apex (Molendijk et al., 2001; Jones et al., 2002; Carol and Dolan, 2006). Recently, a number of receptor-like kinases (RLKs) have been identified that are involved in cellular growth regulatory mechanisms, especially in cell elongation associated with root hair tip growth in higher plants (Shiu and Bleecker, 2001; Lehti-Shiu et al., 2009; Lindner et al., 2012). In particular, CrRLK1L subfamily proteins, which includes *FER* (Huck et al., 2003), *ERELUS* (*ERE*) (Haruta et al., 2014; Schoenaers et al., 2018), *THESEUS1* (*THE1*) (Hematy et al., 2007), *ANXUR1/2* (Miyazaki et al., 2009), have been implicated in cell wall sensing associated with a variety of cellular events such as female fertility, cell elongation, root-hair development, mechanosensing, and responses to hormones and pathogens (Boisson-Dernier et al., 2009; Cheung and Wu, 2011; Lindner et al., 2012).

The CrRLK1L subfamily is named after the first member functionally characterized in *Catharanthus roseus* cell cultures (Schulze-Muth et al., 1996), and Arabidopsis, contains 17 CrRLK1L subfamily members (Hematy and Hofte, 2008). The majority of CrRLK1L receptor-like kinase proteins are predicted serine/threonine kinases with a single transmembrane between an N-terminal extracellular domain with two tandem domains showing limited homology to the carbohydrate-binding domain of animal malectin proteins and a C-terminal cytoplasmic kinase domain (Schallus et al., 2008); (Cheung and Wu, 2011). *THE1* was discovered in a screen for suppressors that partially restored the dark-grown hypocotyl growth defect of *procuste1-1* (*prc1-1*), which is defective in the cellulose synthase catalytic subunit CESA6 (Hematy et al., 2007), and, *HERCULES1* (*HERK1*), was identified as functionally redundant with *THE1* in modulating cell elongation (Guo et al., 2009). While single mutant *the1* or *herk1* plants displayed normal growth, *the1 herk1* double mutants were severely stunted (Guo et al., 2009). *ANXUR1* and *ANXUR2* (*ANX1* and *ANX2*), are exclusively expressed in the male gametophyte (Boisson-Dernier et al., 2009), and function in maintaining pollen tube wall integrity during migration through floral tissues until their deactivation allows the pollen to burst during fertilization (Boisson-Dernier et al., 2009).

Similar to *ANX1* and *ANX2, FER*, which is allelic to *SIRÈNE (SRN)*, was initially identified in the regulation of female control of fertility (Huck et al., 2003), and is highly expressed in the synergid cells of the female gametophyte and in a variety of vegetative tissues, but not in the male gametophyte (Escobar-Restrepo et al., 2007; Guo et al., 2009). In the female gametophyte, *FER* is involved in sensing pollen tube arrival and promoting its rupture, (Huck et al., 2003; Rotman et al., 2003), and in the initiation of programmed cell death of one of two synergid cells during the double fertilization event (Ngo et al., 2014). *FER* also inhibits polyspermy through regulation of demethylesterified pectin accumulation in the filiform apparatus of the ovule (Duan et al., 2020).

In addition to important roles during fertilization, *FER* has also been shown to regulate aspects of root hair elongation (Duan et al., 2010; Huang et al., 2013), calcium signaling during mechanical stimulation of roots (Shih et al., 2014), and cell wall responses to both abiotic and biotic stress (Huck et al., 2003; Rotman et al., 2003; Lindner et al., 2012; Duan et al., 2014; Ngo et al., 2014; Shih et al., 2014; Li et al., 2016). *FER* was identified as a ROP guanidine exchange factor 1 (ROPGEF1) that activates ROP GTPase signaling during root hair tip-growth in Arabidopsis (Duan et al., 2010). *FER*, and other members of the CrRLK1L receptor families have been proposed to bind to secreted RALF (<u>r</u>apid <u>al</u>kalinization <u>f</u>actor) peptide ligands (Haruta et al., 2014), with RALF1 binding the *FER* extracellular domain to suppress cell elongation of the primary root (Haruta et al., 2014). RALF34 binds to *THE1* during regulation of lateral root initiation (Gonneau et al., 2018), and RALF23 binding to *FER* negatively regulates plant immune responses (Stegmann et al., 2017). More recently, *FER* was shown to associate with a GPI-anchored co-receptor, *LORELEI* (*LRE*; Liu et al., 2016), and a crystal structure of the FER extracellular domain bound to a LRE-like protein (LLG2), and RALF23 shows that LLG2 and RALF23 peptides associate with the second malectin-like domain (Xiao et al., 2019).

Although RALF peptide binding occurs in several CrRLK1L receptor-like kinases, it remains unclear whether this is the only ligand recognized by this class of receptor-like kinases. Indeed, *the1-1* mutants displayed altered growth phenotypes despite the *the1-1* mutant extracellular domain being able to bind RALF34 (Gonneau et al., 2018). Additionally, while RALF peptides appear to only bind the second of two malectin-like domains in the FER extracellular domain, both these domains appear to associate with pectin polysaccharides (Feng et al., 2018). We have identified a new temperature-sensitive mutation (*fer-ts*) in a highly conserved glycine residue (G41S) present in the extracellular domain of the FER receptor kinase, as well as other members of the CrRLK1L receptor-like kinase family and mammalian malectin sequences. The *fer-ts* mutant exhibited rapid cessation of root hair tip-growth within one minute upon transferal to elevated (30°C) growth temperatures. This rapid cessation of root hair tip growth was also observed when *fer-4* null mutants were transformed with a fluorescently-tagged version of the temperature-sensitive FER(G41S)-EYFP. FER(G41S)-EYFP was still present in the plasma membrane in root hairs that had ceased tip-growth, indicating that the primary defect of this mutant is due to failure to properly transmit extracellular signals at elevated temperatures. These findings provide a powerful new tool for dissecting how FERONIA, and potentially other CrRLK1L receptor-like kinases, function during plant growth and development.

## RESULTS

### Isolation of a temperature-sensitive mutant that inhibits root hair tip growth

The regulatory GTPase, RabA4b, participates in membrane trafficking associated with the polarized secretion of cell wall components in plant cells. In addition, loss of tip localization of EYFP-RabA4b is highly correlated with inhibition of root hair tip growth (Preuss et al., 2004). In order to identify molecular components that control root hair tip growth, EMS-mutagenized seeds of a stable transgenic Arabidopsis line expressing EYFP-RabA4b were screened for seedlings with wild-type root hairs at normal temperatures (20°C), but which displayed impaired root hair growth when grown at elevated temperatures (30°C). The progeny of approximately 6,000 EMS-mutagenized seeds were screened. From the screening, four temperature-sensitive (ts-) root hair growth defect mutants were isolated, which we initially termed <u>L</u>oss-of-<u>T</u>ip-<u>L</u>ocalization mutants (*ltl1* to *ltl4*). Among these *ltl ts* mutants, *ltl2*, (subsequently referred to as *fer-ts*) root hair growth characteristics were examined under normal and elevated temperature conditions. In normal growth conditions (20°C), *fer-ts* root hairs displayed normal growth, however, both root hair growth and apical accumulation of EYFP-RabA4b of *fer-ts* root hairs were dramatically inhibited at 30°C (Figure 1a and 1b).

**Figure 1.**
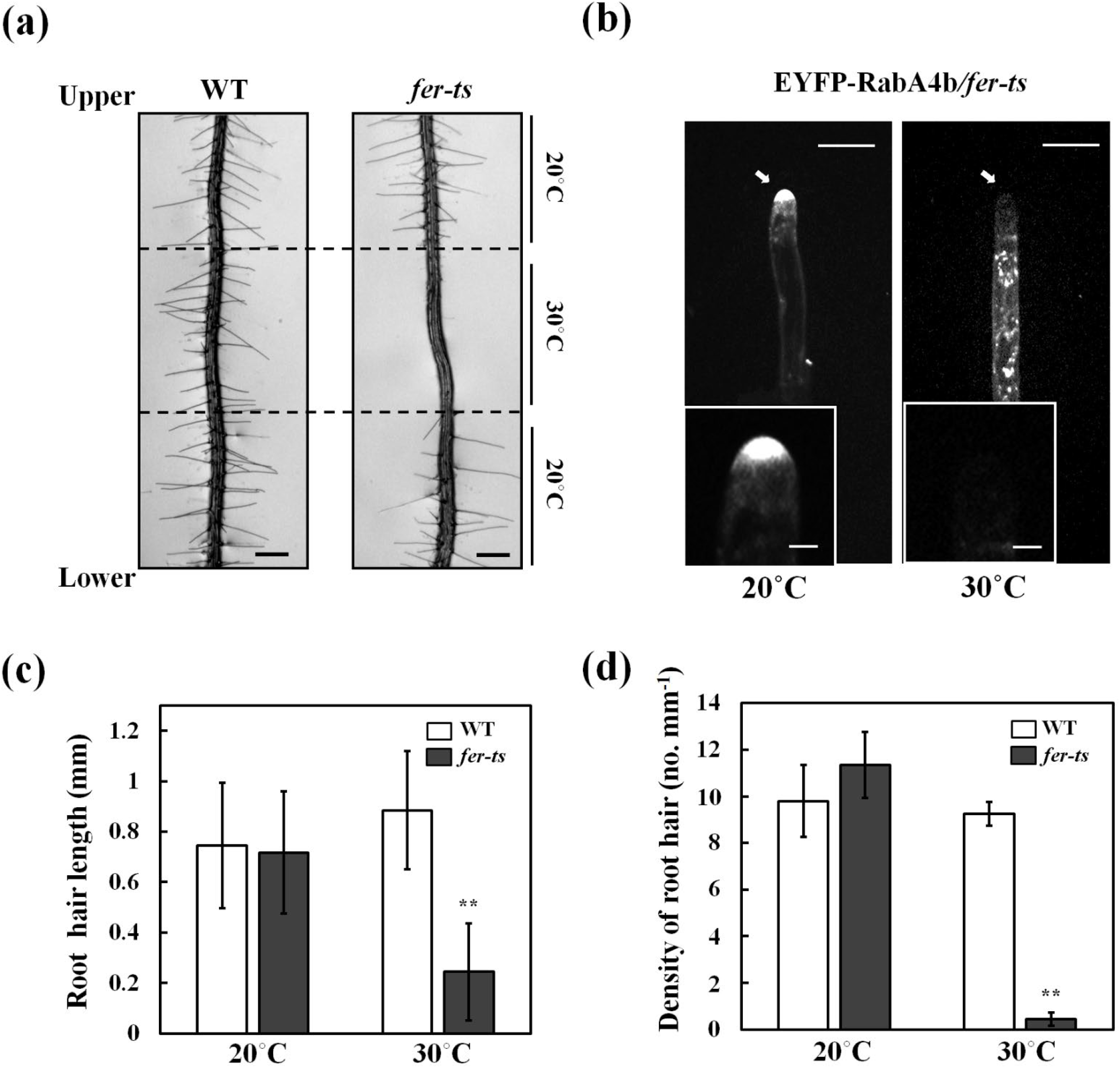
Isolation of a temperature-sensitive root hair growth defect mutant. (a) Seven day-old seedling plants were grown vertically in 1/4 MS media under 20°C and transferred to 30°C for 6 h, followed by 24 h recovery at 20°C. Bright field images were collected with a Nikon Eclipse E600 wide-field microscope with a 10x Plan Apo DIC (0.75 NA) lens. Dashed lines indicate root tip positions when seedling plants were transferred to 30°C condition for 6 hours, and again when they were transferred back to 20°C. Scale bars = 200 μm. (b) Localization of EYFP-RabA4b protein in growing roothair cells of *fer-ts* mutants at 20°C and 30°C. Medial root hair sections were collected using spinning-disk confocal microscopy from growing root-hair cells of seven-day-old seedlings stably expressing EYFP-RabA4b in the *fer-ts* mutant in 20°C (left) or 30°C (right) using a Zeiss 40x Plan-Apochromat (1.3 NA) lens and appropriate EYFP fluorescence filter sets. Scale bars = 10 μm. Insets, magnified images to show details of EYFP-RabA4b subcellular localization in root-hair tips. Scale bars = 2 μm. (c) Quantification of root hair length in WT and *fer-ts* mutants under 20°C (wild-type (n=392), *fer-ts* (n=454)) and 30°C (wild-type (n=185), *fer-ts* (n=23)) conditions. (d) Calculation of root hair densities in WT and *fer-ts* mutants at 20°C (wild-type (n=392), *fer-ts* (n=454)) and 30°C (wild-type (n=185), *fer-ts* (n=23)) in fully expanded primary roots of seven-day old plants. In each case, root hair lengths and densities were measured from n=20 individual seedlings. Error bars represent SD. **p<0.001 by Student’s *t*-test.

To quantify root hair elongation in *fer-ts* under normal and elevated temperature conditions, both root hair lengths and root hair density were measured. No significant differences were found between wild-type and *fer-ts* either in mature root hair length, or in the number of root hairs per unit root length when plants were grown at 20°C. However, both length of root hairs and root hair density were greatly reduced in *fer-ts* in plants grown at 30°C (Figure 1c and 1d). Primary root length and root growth rates of *fer-ts* seedlings were indistinguishable from wild-type at 20°C, but while wild-type roots grew slightly faster at 30°C, *fer-ts* primary root growth rates remained unchanged at elevated temperatures (See Supplemental Figure S1). These results indicated that, at least in early stages of seedling growth and development*fer-ts* temperature sensitive defects are largely specific to root hair elongation in elevated temperature conditions.

In order to characterize effects of the *fer-ts* mutation on root hair growth dynamics, elongating root hairs were visualized by time-lapse microscopy for two hours. Seven day-old *fer-ts* seedlings were placed in a temperature-controlled plant growth chamber at 20°C for 50 min and then the temperature of the chamber was rapidly transitioned to 30°C (Figure 2). While root hair growth was unaffected by temperature transition in wild-type plants, transition from normal to elevated temperatures resulted in rapid cessation of tip-growth in the *fer-ts* mutant (Figure 2 and Movie S1).

**Figure 2.**
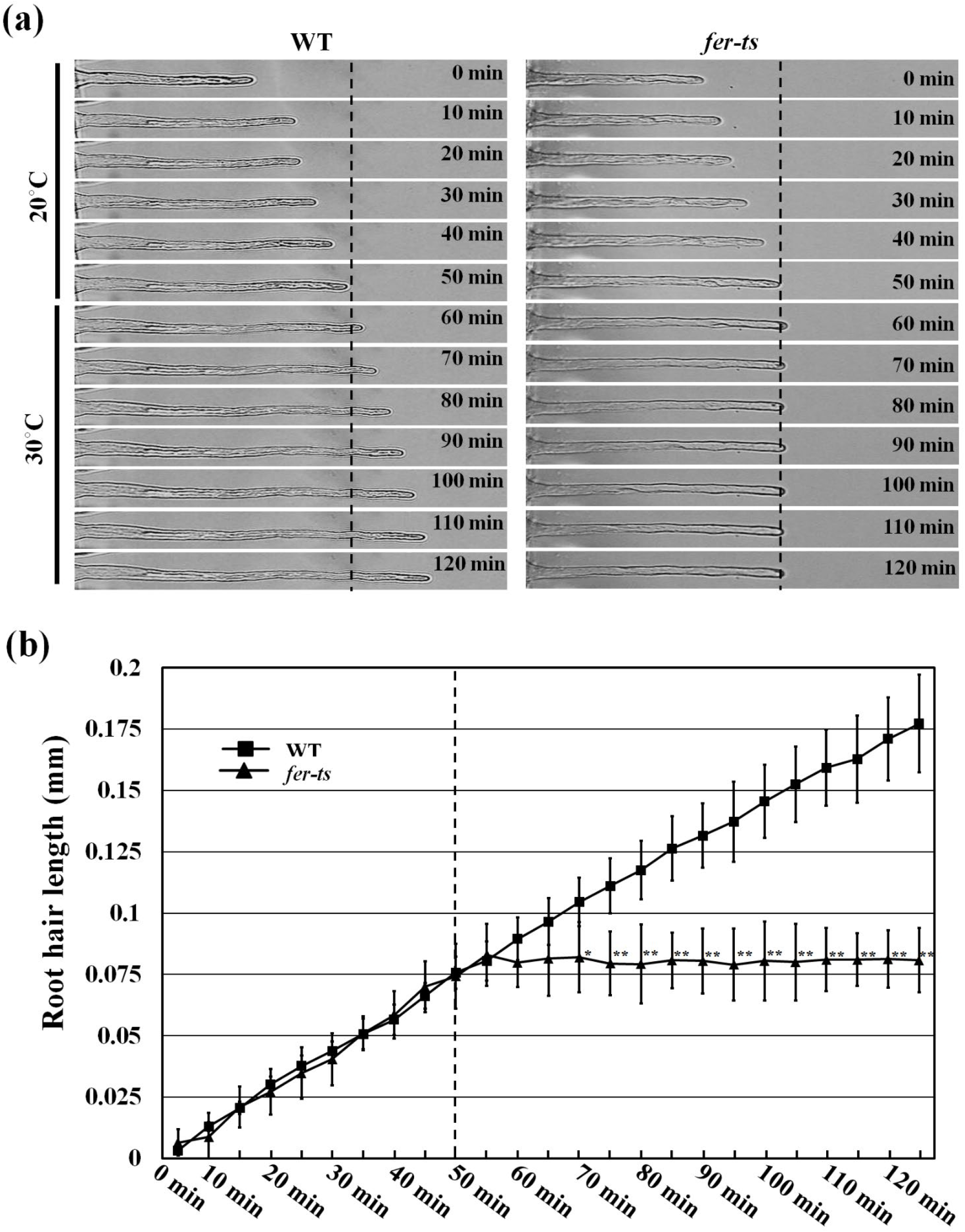
Root hair growth dynamics in WT and *fer-ts* seedlings. (a) Root hair tip-growth in WT and *fer-ts* mutant plants under normal (20°C) and elevated (30°C) temperatures by time-lapse microscopy. Bright field images of growing root hairs of WT and *fer-ts* mutant plants were collected every minute by time-lapse microscopy using a Zeiss 40x Plan-Apochromat (1.3 NA) lens. Representative images of WT and *fer-ts* mutant root hair elongation are presented at 10 min intervals. (b) Quantitative analysis of WT (n=4) and *fer-ts* mutant (n=4) root hair lengths upon transition to 30°C. WT (black squares) and *fer-ts* (black triangles) mutant root hair elongation were measured every five minutes and root hair lengths were determined using the measure function in Image J. The dashed line indicates transition from 20°C to 30°C. Error bars represent SD. *p<0.05, **p<0.01 by Student’s *t*-test.

Since root hair growth is tightly linked to accumulation of EYFP-RabA4b compartments in root hair tips (Preuss et al., 2004; Preuss et al., 2006; Thole et al., 2008), we were interested how this localization was affected upon transition from 20°C to 30°C in the *fer-ts* mutants. EYFP-RabA4b was detected in the apical region of growing *fer-ts* root hairs at 20°C, but tip localization of EYFP-RabA4b was rapidly lost upon transition to 30°C (Figure 3). EYFP-RabA4b accumulation was examined in wild-type and *fer-ts* plants in a temperature-controlled chamber at 20°C for 14 min, and then the chamber was rapidly transitioned to 30°C (Figure 3a and Supplemental Figure S2). Images of growing root hairs were collected at one minute intervals by time-lapse confocal microscopy, and tip-localized EYFP signal was quantified in both wild-type (Figure 3b) and *fer-ts* (Figure 3c) root hairs. While tip-localized EYFP-RabA4b signal was unaffected by transition from 20°C to 30°C in wild-type root hairs (Figure 3b), tip-localized EYFP-RabA4b was significantly reduced in *fer-ts* within one minute of the transition from 20°C to 30°C (Figure 3c).

**Figure 3.**
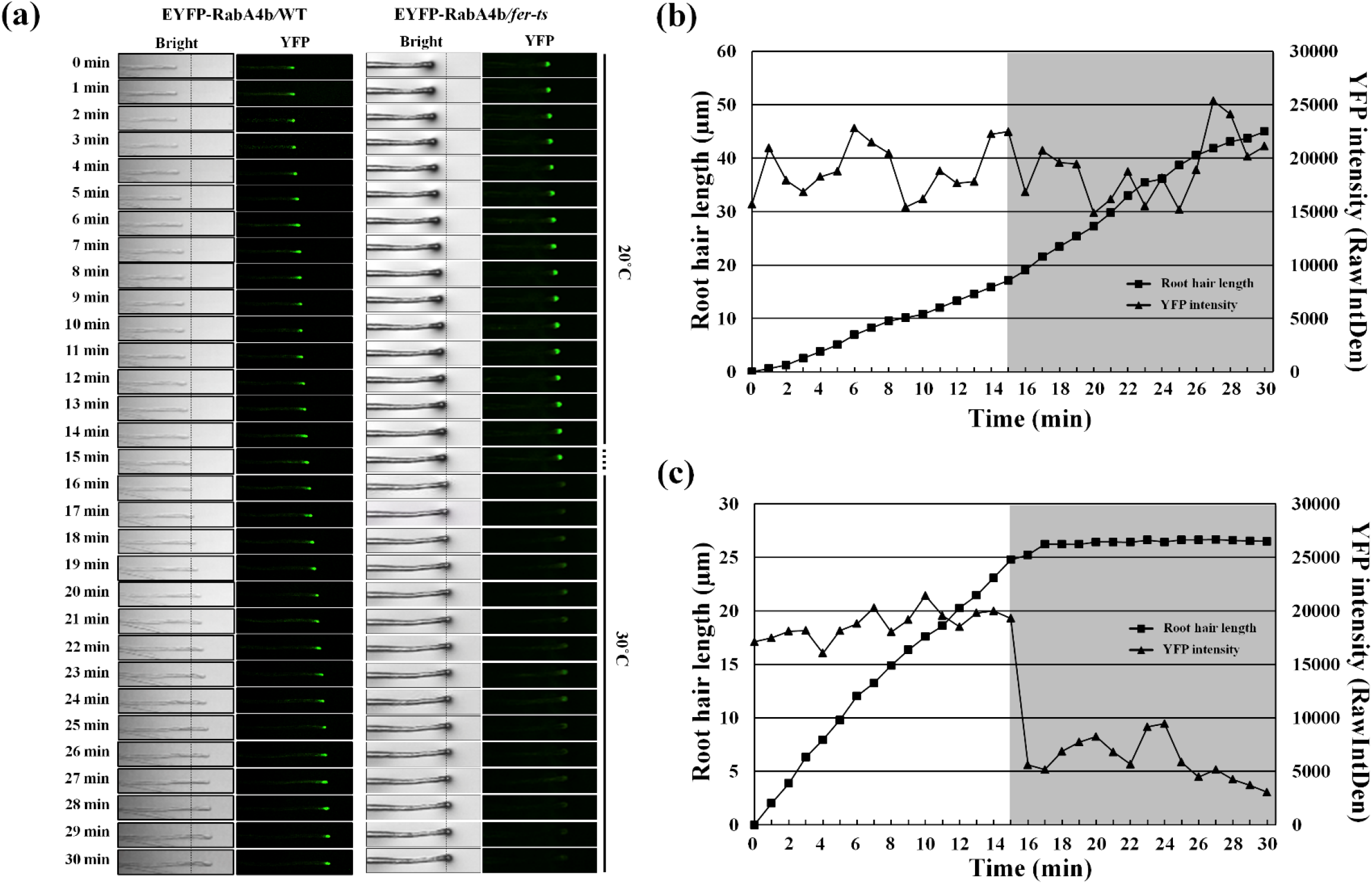
Temperature-sensitive subcellular dynamics of EYFP-RabA4b labeled compartments in growing root hairs in *fer-ts* mutants. (a) Localization of EYFP-RabA4b protein in a growing root-hair cells of wild-type and *fer-ts* mutant plants at normal (20°C) or elevated (30°C) temperature conditions. Growing wild-type and *fer-ts* root hairs were imaged at one minute intervals for 30 min at 20°C at which the growth chamber temperature was raised to 30°C. For each time point, both bright field (Bright) and fluorescence (YFP) images were collected sequentially, and tip-localized EYFP-RabA4b compartments were monitored by spinning-disk fluorescence confocal microscopy using a Zeiss 40x Plan-Apochromat (1.3 NA) lens with appropriate EYFP fluorescence filter sets. The grey box indicates when the temperature was raised to 30°C. Quantification of root hair elongation and EYFP-RabA4b root hair tip localization in wild-type (b) and *fer-ts* mutant (c) plants. Root hair length and EYFP-RabA4b fluorescence were quantified by Image J every 1 min.

### Map-based cloning and full-genome sequencing of the *fer-ts* locus

To identify the mutant locus responsible for the rapid, temperature-sensitive loss of root hair elongation and tip-localized EYFP-RabA4b, map-based cloning and full-genome sequencing was performed. F_2_ mapping populations were obtained by reciprocal crosses of back-crossed mutants (Col-0) with L*er* wild-type plants (Bell and Ecker, 1994). Segregating F_2_ populations were used for the subsequent map-based cloning. The temperature-sensitive mutant lesion was mapped to chromosome 3 between the SSLP markers *NIT1.2* and *NGA6* (Figure 4a). Subsequent low-resolution mapping and whole genome sequencing identified that the *fer-ts* mutant locus was due to a G121A nucleotide replacement resulting in a G41S substitution mutation within the extracellular domain of the previously characterized FER receptor-like kinase (Figure 4b and 4c). In order to eliminate the possibility that the G➔A substitution that gave rise to the *fer-ts* G41S mutation influenced accumulation of *FER* mRNA at the transcriptional level, we performed RT-PCR analysis. *FERONIA* transcript levels were unchanged from those in wild-type plants (Figure 4d).

**Figure 4.**
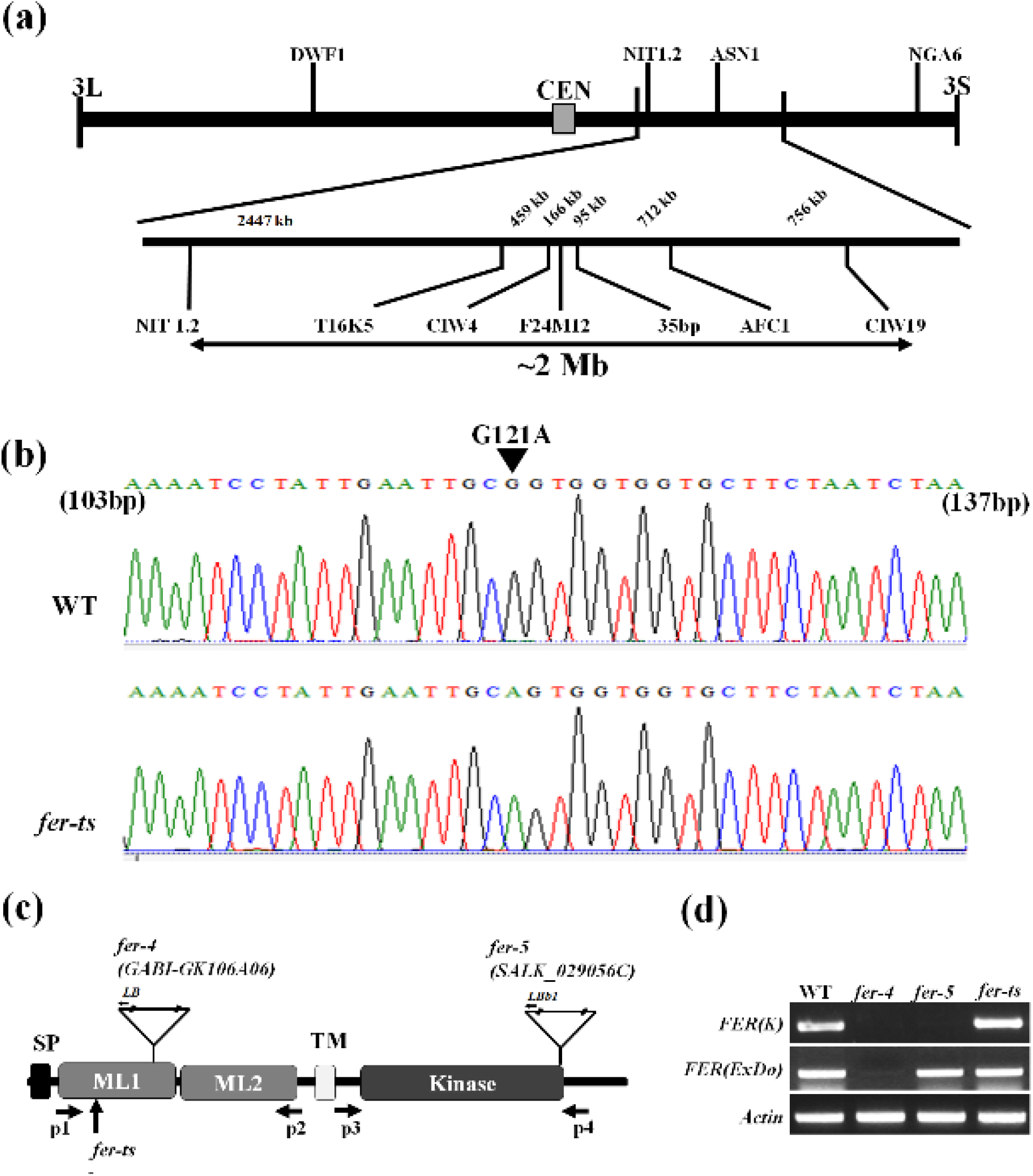
Map-based cloning of *fer-ts*. (a) A linear diagram of the Arabidopsis third chromosome is shown, with a magnified F24M12 marker region displayed below. The centromere is indicated with filled-rectangle. Low-resolution map-based cloning resulted in identification of the *fer-ts* locus within an approximately 2 Mb region of chromosome III bounded by markers NIT1.2 and CIW19 (Double headed arrow). (b) SNPs specific to the ts-mutant within this region were identified using whole genomic resequencing, followed by targeted resequencing of genomic DNA sequencing of *fer-ts* and wild-type parental lines. A single G➔A substitution was found in *FERONIA* (At3g51550). The arrowhead indicates G121A substituted mutation in *FERONIA* gene locus. (c) Schematic diagram of FERONIA protein domains and mutation regions, composed of an N-terminal extracellular domain (tandem repeat malectin-like domains; ML1 and ML2), TM (transmembrane) domain in the middle region and a C-terminal kinase domain (serine/threonine kinase), end of N-terminus has signal peptide (SP) sequence for plasma-membrane trafficking. The *fer-4* and *fer-5* mutants displayed that T-DNA was inserted in malectin-like domain 1 and kinase domain, respectively. (d) RT-PCR analysis of T-DNA inserted mutants and EMS mutants. The FER(K) and FER(ExDo) domains were amplified using P1 and P2 primers and P3 and P4 primers, respectively. Actin was used as a loading control.

### The G41S mutation in *fer-ts* is a highly conserved glycine residue in CrRLK1L subfamily proteins

The G41S substitution of the *fer-ts* mutant rapidly inactivated *FER* function during root hair growth at elevated temperatures. In addition, a similar G37D mutation is responsible for inactivation of THE1, another member of the CrRLK1L family (Hematy et al., 2007), and multiple sequence alignment analysis with other Arabidopsis CrRLK1L family members and animal malectin sequences showed that the G41 residue of FERONIA is absolutely conserved in these malectins and malectin-like 1 (ML1) domains (Figure 5a, and Supplemental Figure S3). Interestingly, based on structural studies of animal malectin proteins, five key residues (Y67, Y89, Y116, F117, D186; Figure 10b, red residues) were found to form contacts with a bound disaccharide ligand, nigerose, in the active site as determined by structural analysis of the *X. laevis* malectin protein (Schallus et al., 2008; Muller et al., 2010). In this malectin structure these surface exposed residues extend from the malectin fold forming the nigerose binding pocket, with the conserved glycine (G40) the bottom of this structural region (Figure 5b). While several of the tyrosine and phenylalanine residues shown to be important for interaction with carbohydrates in animal malectin proteins are maintained in plant malectin-like domains (e.g. FERONIA Y88, Y114, F115, D197) (Figure 5a, and Supplemental Figure S3), these are not surface exposed in the ML1 domain of the recently described crystal structure of FER (Figure 5c, in green) with its co-receptor LLG2 (Figure 5b, in blue) and a RALF23 ligand (Figure 5c, in magenta) (Xiao et al., 2019). It is however notable that in this structure the invariant glycine (G41; Figure 5c, red residue) of the FER ML1 domain is not located within or nearby the RALF23 and LLG2 binding surfaces in the ML2 domain (Xiao et al., 2019).

**Figure 5.**
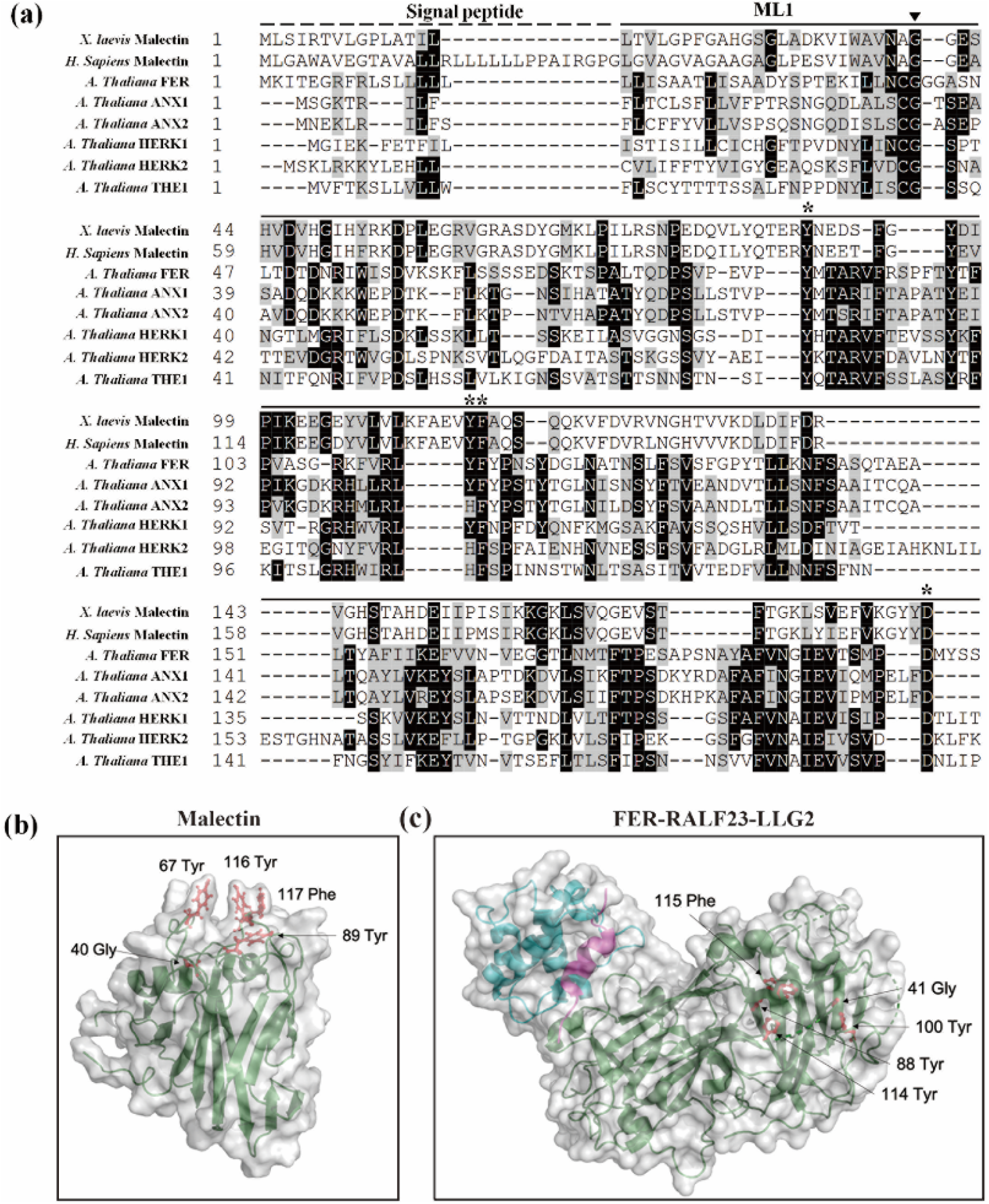
Localization of the highly conserved glycine mutation in fer-ts in *A. thaliana* FERONIA:RALF23:LLG2 and *X. laevis* malectin structures. (a) Several residues important for binding carbohydrate ligands are conserved in plant CrRLK1L receptor kinase family members. Sequences analyzed include animal Malectin (*X. laevis* and *H. sapiens*) FERONIA and other well-characterized CrRLK1L homologs in *A. thaliana* (ANXUR1; ANX1, ANXUR1; ANX2, HERCULES1; HERK1 and THESEUS1; THE1). Putative N-terminal signal peptides are indicated as black dashed lines, and malectin and CrRLK1L ML1 domains by solid lines, respectively. The highly conserved G41 of FER is marked by arrowhead. Black boxes indicate fully conserved residues; shaded boxes indicate similar and partially conserved residues. Conserved residues that have been shown to participate in binding nigerose in the *X. laevis* malectin structure are marked by asterisks. Sequence alignment analysis was performed by CLUSTAL Omega program (http://www.ebi.ac.uk/Tools/msa/clustalo/) and displayed by using BOXSHADE software (www.ch.embnet.org/software/BOX_form.html). (b) Crystal structure of the *X. laevis* malectin protein (PDB ID: 2K46) with binding pocket aromatic residues and the highly conserved glycine residue based on sequence similarity to FER shown in red. (c) Crystal structure of FER protein (green) in complex with RALF23 ligand (magenta) and GPI-anchored protein LLG2 (blue) (PDB ID: 6A5E). No analogous binding pocket is observed on the ML1 domain, as all conserved aromatic residues (red) are buried within the protein. Both (b and c) were generated using PyMol (DeLano Scientific).

### *fer-ts* phenotypes were confirmed by reciprocal crossing with FERONIA mutants and complementation assays

To confirm that the temperature sensitive root hair defects and loss of tip-localized EYFP-RabA4b were causally linked to the G41S mutation in the FER locus, *fer-ts* mutant plants were reciprocally crossed with two previously characterized *FER* mutants, *fer-4* and *fer-5* (Duan et al., 2010). While both *fer-4* and *fer-5* fail to accumulate root hairs, *fer-4* was shown to fully abolish FER protein, while *fer-5* was shown to accumulate a truncated FER protein missing a functional cytosolic protein kinase domain (Duan et al., 2010). As a control, F1 plants of *fer-ts* crossed to wild-type plants displayed normal root hair growth in both normal and elevated temperature conditions, confirming that the *fer-ts* mutation is recessive. However, F1 plants of either *fer-ts* (paternal line) crossed with *fer-4* and *fer-5* mutants (maternal lines; Figure 6a), or *fer-ts* (maternal line) crossed with *fer-4* and *fer-5* mutants (paternal line; Supplemental Figure S4a) displayed ts-phenotypes at elevated temperatures, respectively. These F1 plants were confirmed by genomic DNA PCR analysis with primers that discriminated between the *fer-ts* (or wild-type) *FER* loci and *fer-4* and *fer-5* T-DNA insertion mutant loci (Figure 6b-c, and Supplemental Figure S4b). To verify that the FER G41S mutation specifically conferred the temperature-sensitive root hair phenotype, a fluorescently tagged FER-EYFP containing the G41S mutation, FER(G41S), driven by endogenous *FER* promoter sequences, was transformed into *fer-4* and *fer-5* mutant plants. Transgenic *fer-4* and *fer-5* plants, expressing mutant FER(G41S)-EYFP proteins rescued root hair growth defects in these two *fer* mutant backgrounds in a temperature-sensitive manner (Figure 6d). Finally, a wild-type fluorescently-tagged FER-EYFP, driven by endogenous FER promoter sequences, was able to fully rescue *fer-ts* root hair defects. FER(WT)-EYFP protein was detected in plasma membranes and *fer-ts* transgenic lines stably transformed and homozygous for FER(WT)-EYFP displayed the normal root hair growth both 20°C and 30°C (Figure 6e). As observed previously (Duan et al., 2010), pFER-FER(WT)-EYFP fluorescent fusions localized primarily to plasma membranes in both root cortical and mature root hair cells in these transgenic lines, although a small amount of pFER-FER(WT)-EYFP could also be seen in subcellular compartments that colocalized with FM 4-64 (Figure 6f) Taken together, these data strongly support that *fer-ts* phenotype is the result of the G41S mutation of the FERONIA protein.

**Figure 6.**
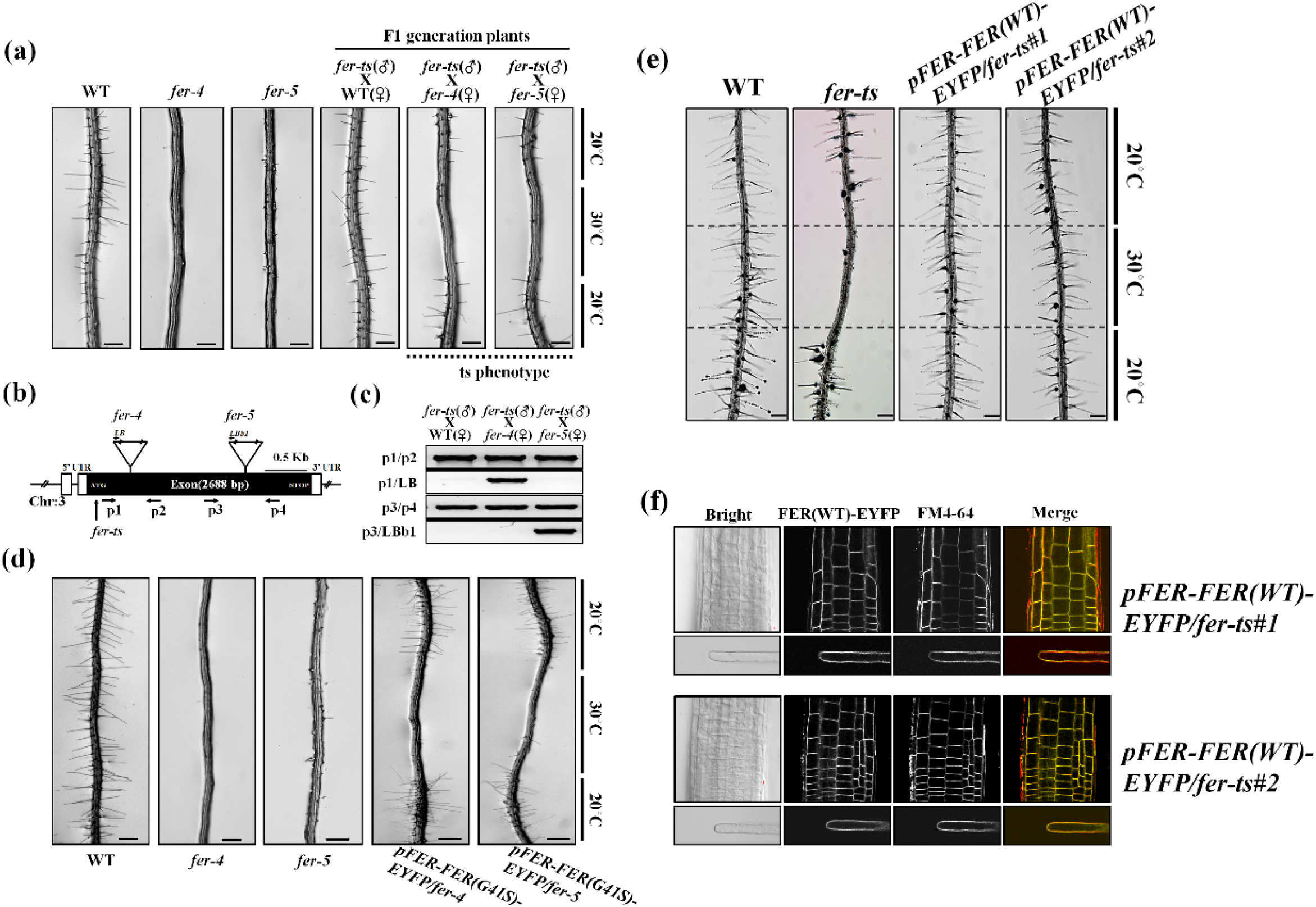
The *fer-ts* mutant confers ts-root hair growth defects when crossed with *fer-4* and *fer-5* mutants. (a) Wild-type (WT), *fer-4, fer-5*, and F1 progeny from crosses (paternal = *fer-ts*, maternal = *fer-4* or *fer-5*) of *fer-ts/fer-4* and *fer-ts/fer-5* were grown vertically for seven days at 20°C, transferred to 30°C for 6 h, and then grown for an additional 24 h at 20°C. Both *fer-ts/fer-4* and *fer-ts/fer-5* progeny clearly demonstrated a ts-dependent root hair phenotype. Scale bars = 200 μm. (b) Schematic diagram of the *FERONIA* gene structure. Open and filled boxes indicate untranslated regions (UTRs) and exon regions, respectively. The locations of T-DNA insertion mutants (*fer-4* and *fer-5*) and *fer-ts* are indicated by triangles and arrows, respectively. (c) Genotyping of crossed F1 plants. Genomic DNA was extracted from F1 generation plants and subjected PCR to confirm presence of the *fer-4* and *fer-5* genotypes (d) Both *fer-4* and *fer-5* display temperature-dependent root hair phenotypes when transformed with a fluorescently-tagged FER construct containing the *fer-ts* mutation (pFER-FER(G41S)-EYFP). Seven-day old seedlings stably transformed and homozygous for (pFER-EYFP(G41S)-EYFP) were grown vertically at 20°C, transferred to 30°C for 6 h, and then grown for an additional 24 h at 20°C. Presence of the transgenic pFER-FER(G41S)-EYFP construct clearly demonstrated a ts-dependent root hair phenotype. Scale bars = 200 μm. (e) Temperature sensitive root hair growth defects of *fer-ts* are fully rescued when transformed with a fluorescently-tagged FER construct (pFER-FER(WT)-EYFP). Seven-day old seedlings stably transformed and homozygous for (pFER-EYFP(WT)-EYFP)were grown vertically at 20°C, transferred to 30°C for 6 h, and then grown for an additional 24 h at 20°C. Bright field images were collected with a Nikon Eclipse E600 wide-field microscope with a 10x Plan Apo DIC (0.75 NA) lens. (f) Subcellular localization of FER(WT)-EYFP protein in roots and root hairs in a rescued *fer-ts* mutant plant. Fluorescent confocal images displaying the subcellular distribution of FER(WT)-EYFP protein was detected from growing root, and mature root hair cells of seven-day-old *fer-ts* seedlings stably transformed and homozygous for pFER-FER(WT)-EYFP. Cells were counter-stained by incubating for 5 min in a FM4-64 to visualize cell membranes. Images were collected by spinning-disk fluorescence confocal microscopy using a Zeiss 40x Plan-Apochromat (1.3 NA) lens with appropriate EYFP and FM4-64 fluorescence filter sets. Scale bars = 20 μm.

### FERONIA is primarily localized to the plasma membranes and the *fer-ts* G41S substitution does not alter its subcellular localization

Previously, GFP-fused FERONIA was shown to localize to plasma membranes in various plant tissues (Duan et al., 2010). At normal temperatures, FER(WT)-EYFP was observed primarily in plasma membranes in various tissues such as leaf, root and in mature root hairs (Figure 6f and 7a), and at higher magnification in root cortical cells FER(WT)-EYFP was restricted to plasma membranes, and did not display significant accumulation in intracellular compartments in non-tip-growing root cortical cells (Figure 7b). Interestingly, in actively growing root hairs, FER(WT)-EYFP protein was observed both in plasma membranes and an apical localized vesicle population in growing root hairs (Movie S2). To determine whether the introduction of the G41S substitution in the *fer-ts* mutant might affect the subcellular localization of this protein in normal or elevated temperatures, we compared the subcellular distributions of FER(WT)-EYFP and FER(G41S)-EYFP at both 20°C and 30°C (Figure 7c and 7d). While no changes in accumulation or distribution of FER(WT)-EYFP were observed in roots and mature root hairs between 20°C and 30°C conditions (Figure 7c), at 30°C some FER(G41S)-EYFP fluorescence could be observed in internal subcellular membranes, although significant levels of the FER(G41S)-EYFP remained at the plasma membranes in these cells even after incubation at 30°C for 6 hours (Figure 7e).

**Figure 7.**
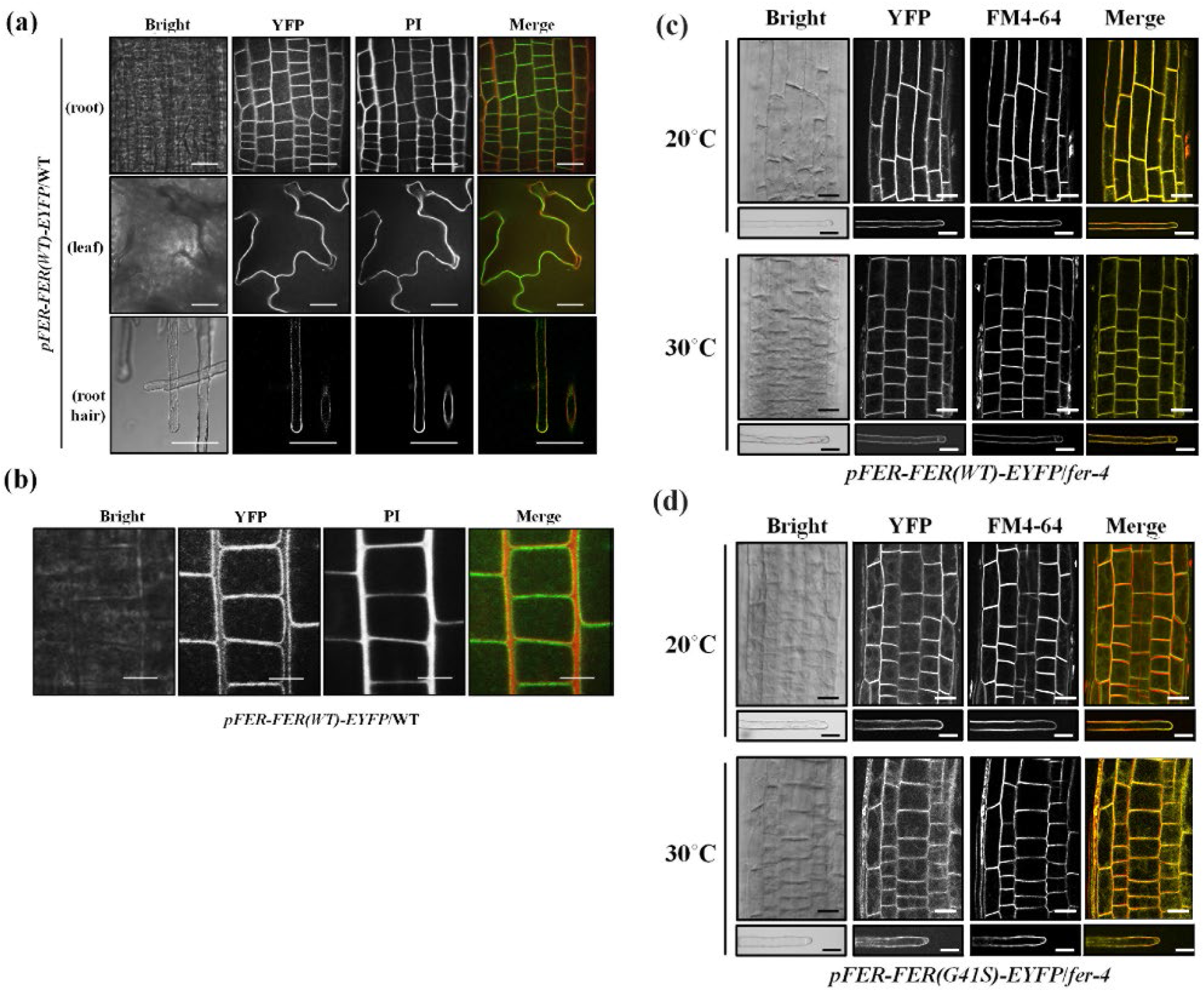
Subcellular localization of FER(WT)-EYFP and FER(G41S)-EYFP fluorescent fusion proteins in stably transformed *Arabidopsis*. (a) Subcellular localization of FER(WT)-EYFP protein in various tissues. Fluorescent confocal images displaying the subcellular distribution of FER(WT)-EYFP protein was detected from growing root, leaf and root hair cells of seven-day-old seedlings in *pFER-FER(WT)-EYFP*/WT transgenic plants. Cell walls were counter-stained by incubating for 5 min in a propidium iodide (PI) solution (10 μg/ml). Images were collected by spinning-disk fluorescence confocal microscopy using a Zeiss 40x Plan-Apochromat (1.3 NA) lens with appropriate EYFP and PI fluorescence filter sets. Scale bars = 20 μm. (b) Magnified images of FER(WT)-EYFP fluorescence. FER(WT)-EYFP in wild-type of growing root cells of seven-day-old *A. thaliana* seedlings was detected by spinning-disc confocal microscopy using a Zeiss 100x Plan-Apochromat (1.46 NA) oil immersion objective with appropriate EYFP and PI filter sets. Scale bars = 10 μm. (c-d) Subcellular localization of FER(WT)-EYFP (c) and FER(G41S)-EYFP (d) fluorescent fusions in root and root hair cells at normal (20°C) and elevated (30°C) temperatures. Cells from *fer-4* plants stably transformed and homozygous for FER(WT)-EYFP or FER(G41S)-EYFP were counterstained with FM4-64 (5 μM) for five minutes to visualize cell membranes. Images were collected by spinning-disk fluorescence confocal microscopy using a Zeiss 40x Plan-Apochromat (1.3 NA) lens with appropriate EYFP and FM4-64 fluorescence filter sets. Scale bars = 20 μm; root, 10 μm; root hair.

Since the *fer-ts* phenotype rapidly inhibited tip-growth upon transition to elevated temperatures (30°C), we were interested to determine how this might effect FER(G41S)-EYFP localization in actively growing root hair cells. Seven day-old *fer-ts* seedlings were placed in a temperature-controlled plant growth chamber at 20°C and images of growing root hairs of plants stably expressing FER(WT)-EYFP or FER(G41S)-EYFP were collected every five seconds for at least five minutes, and then the temperature of the chamber was rapidly transitioned to 30°C (Figure 8a) to observe the effects of elevated temperatures on both root hair elongation as well as subcellular dynamics of FER(WT)-EYFP and FER(G41S)-EYFP proteins. As noted previously, both EYFP(WT)-EYFP and FER(G41S)-EYFP localized to both plasma membranes and apically-localized vesicle populations in growing root hairs at 20°C (Figure 8a and Movie S3). However, while root hair growth and FER(WT)-EYFP localization were unaffected by elevated temperature (30°C), both tip-growth and accumulation of apically-localized vesicles were inhibited within one minute of transition to elevated temperature (Figure 8c, 8d, and Movie S3). While FER(G41S)-EYFP in apically-localized vesicles was rapidly lost upon cessation of tip-growth, plasma membrane localized FER(G41S)-EYFP levels were maintained at similar levels for several minutes at elevated temperatures. Interestingly, after several minutes of elevated temperature the ratio of FER(G41S)-EYFP localized in an apical plasma membrane domain (Figure 8b, red) versus in flanking plasma membrane regions (Figure 8b, blue) declined (Figure 8e), perhaps reflecting either inhibition of secretion to this apical domain, or possibly due to destabilization of the FER(G41S)-EYFP protein at elevated temperatures.

**Figure 8.**
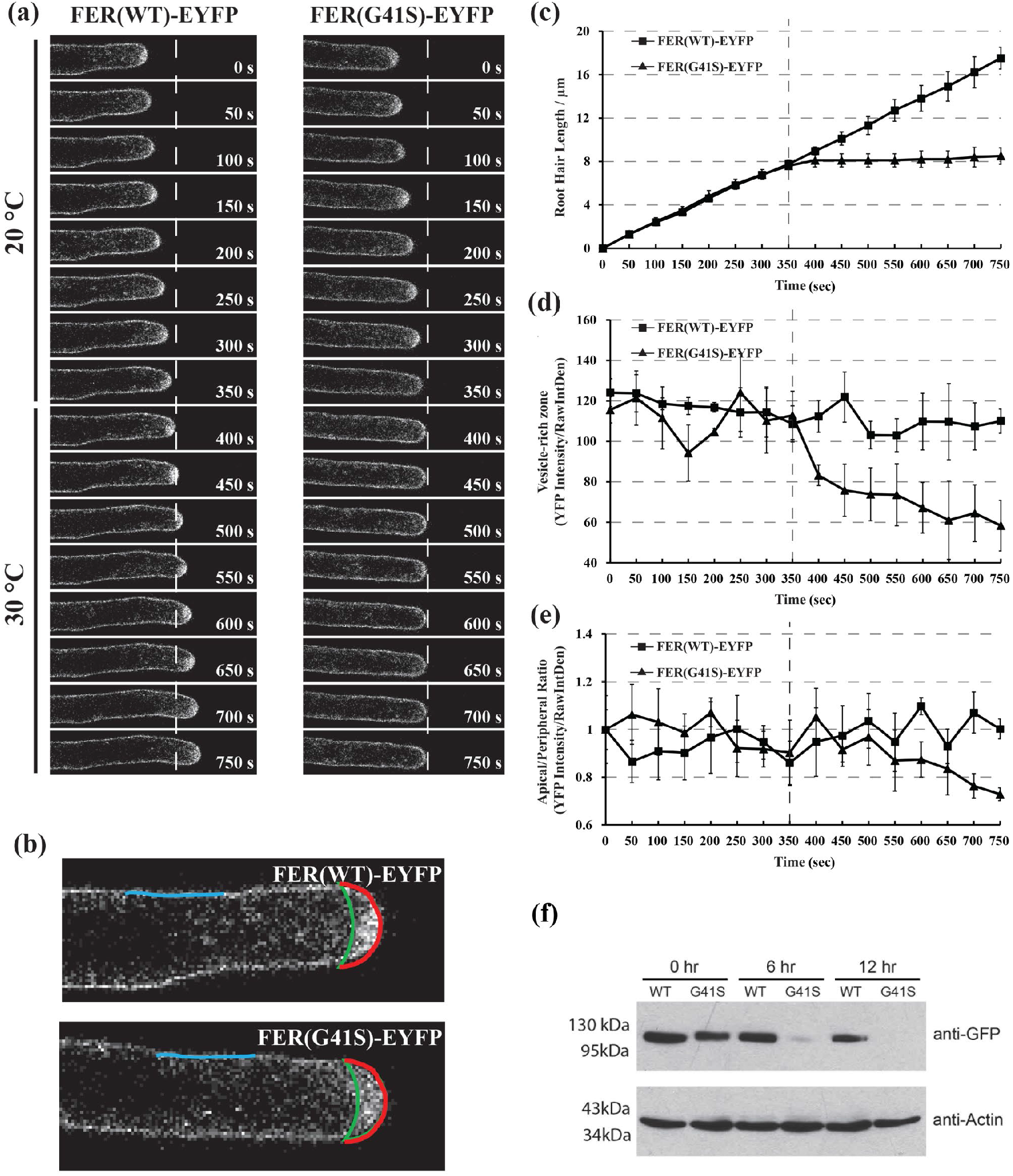
Subcellular localization dynamics of FER(WT)-EYFP and FER(G41S)-EYFP proteins in growing root hair cells under both normal (20°C) and elevated (30°C) temperatures. (a) Root hair tip-growth in stably transformed lines expressing FER(WT)-EYFP and FER(G41S)-EYFP fluorescent fusion proteins under (20°C) and (30°C) temperature by time-lapse microscopy. Confocal images of were acquired using a Leica confocal laser-scanning microscope SP8 with a 63x oil lens (Numerical Aperture = 1.4) with 5s intervals. Representative images are presented at 50s intervals (10 frames). Scale bar: 10 μm. (b) Schematic diagram shown areas and lines used for fluorescence intensity quantification. Apical (red line) and peripheral (blue line) plasma membrane domains were measured, and the area between the apical plasma membrane and green line represents the vesicle-rich zone. (c) Quantitative analysis of root hair growth in stably transformed lines expressing FER(WT)-EYFP (n=3) and FER(G41S)-EYFP (n=3). Root hair elongation was measured every 50s using the measurement function in ImageJ. (d) Quantitative analysis of YFP intensity of vesicle-rich zone. (e) Quantitative analysis of YFP intensity of apical/ peripheral ratio. Values are normalized using 0s as 100%. The dashed line indicates time point of transition from permissive (20°C) to (30°C). Error bars in (c, d, e) represent SD. (f) Protein turnover rates of FER(WT)-EYFP and FER(G41S)-EYFP at elevated temperature (30°C). Five-day old seedling were grown at 20°C and then treated with 200 μM cycloheximide and transferred to 30°C. Total proteins were extracted at each time point and the relative levels were determined using immunoblotting with anti-GFP and anti-actin antibodies. FER(G41S)-EYFP levels rapidly decreased during the time course, while levels of FER(WT)-EYFP were not significantly reduced. Actin was used as a loading control.

In order to examine whether the *fer-ts* mutant protein might display increased protein turnover we blocked new protein synthesis by treating five-day-old *Arabidopsis* seedlings with cycloheximide, and then compared protein turnover rates of the FER(WT)-EYFP and FER(G41S)-EYFP proteins when grown at 30°C. Overall accumulation of the FER(G41S)-EYFP was reduced significantly during the time course, but no significant reduction in protein accumulation was observed for either EYFP(WT)-EYFP or an actin loading control (Figure 8f). These results suggest that while rapid cessation of tip-growth in root hairs occurs within one minute of transition to elevated temperatures, that upon continuous incubation at elevated temperatures *fer-ts* mutant proteins also appear to be subject to enhanced protein turnover.

### The *fer-ts* displays impaired sensitivity to RALF1 peptides in elevated temperature conditions

Signaling in the CrRLK1L family of receptor kinases have been linked to a family of small extracellular peptide hormones called <u>r</u>apid <u>al</u>kalinization <u>f</u>actors (RALFs; (Haruta et al., 2014; Stegmann et al., 2017). RALF1, which was previously demonstrated to suppress cell elongation of the primary root in Arabidopsis and other plants (Pearce et al., 2001), has now been shown to directly bind with the FER extracellular domain (Haruta et al., 2014; Stegmann et al., 2017). Because the G41S mutation in *fer-ts* appears to rapidly inhibit tip-restricted growth at elevated temperatures, perhaps by destabilizing the structure of the extracellular domain of this protein (Figure 8f), we were curious what effect RALF1 peptide might have on root hair growth, and whether the *fer-ts* mutant may display altered responses to RALF1 peptide treatments at elevated temperatures (30°C). Treatment of growing *fer-ts* and wild-type (Figure 9a and Movie S4) root hairs resulted in rapid cessation of tip-growth (<1 min; Figure 9b) in normal temperatures (20°C).

**Figure 9.**
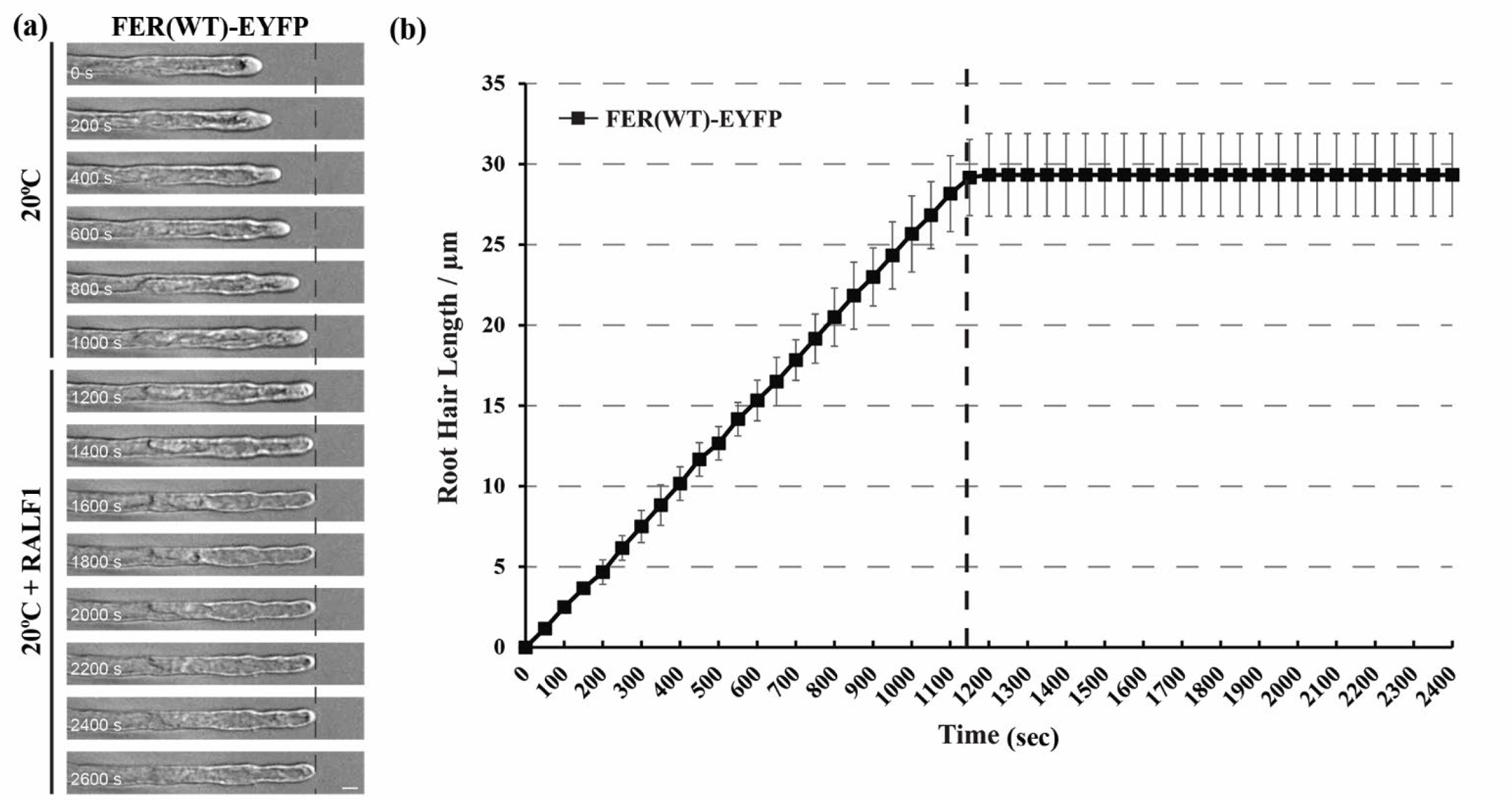
RALF peptide treatment terminates root hair growth in FER(WT)-EYFP expressing line. (a) Root hair tip-growth in stably transformed lines expressing FER(WT)-EYFP fluorescent fusion protein under permissive (20°C) temperature by time-lapse microscopy. Bright field images of growing root hair cells (n=3) were collected using a Nikon Eclipse E600 wide-field microscope with a x 20 Plan Apo (0.45 NA) lens with 5s intervals. The dashed line represents when media containing RALF1 (1 μM) was perfused into the growth chamber. Representative images are presented at 200s intervals. Scale bar: 50 μm. (b) Quantification of root hair growth. Root hair elongation was measured every 50s using the measurement function in ImageJ. Error bars represent SD.

Because *fer-ts* root hair growth was already inhibited at 30°C it was not possible to measure RALF1 peptide effects in root hair cells. However, since primary root elongation did not appear to be as dramatically affected at elevated temperatures as root hair growth (30°C; Figure S1), we examined the temperature-dependent effect of RALF1 peptide treatment on these tissues. As shown in Figure 10, both wild-type seedlings and *fer-ts* mutants were highly sensitive to active RALF1 peptide under normal temperature conditions (20°C). However, as previously described, the sensitivity of root growth to RALF 1 in the *fer-5* mutant was reduced in comparison to wild-type plants at 20°C (Figure 10a and 10c). Importantly, sensitivity of *fer-ts* seedlings to RALF1 peptide treatment was dramatically reduced at 30°C, even though wild-type plants and *fer-5* mutants still responded to RALF1 peptide treatment with similar levels of root elongation inhibition (Figure 10b and 10d). These results support the previous determination that RALF1 peptide signaling occurs through the FER receptor-like kinase, and would be consistent with the G41S mutation resulting in temperature-sensitive inhibition of RALF1-mediated FERONIA signaling.

**Figure 10.**
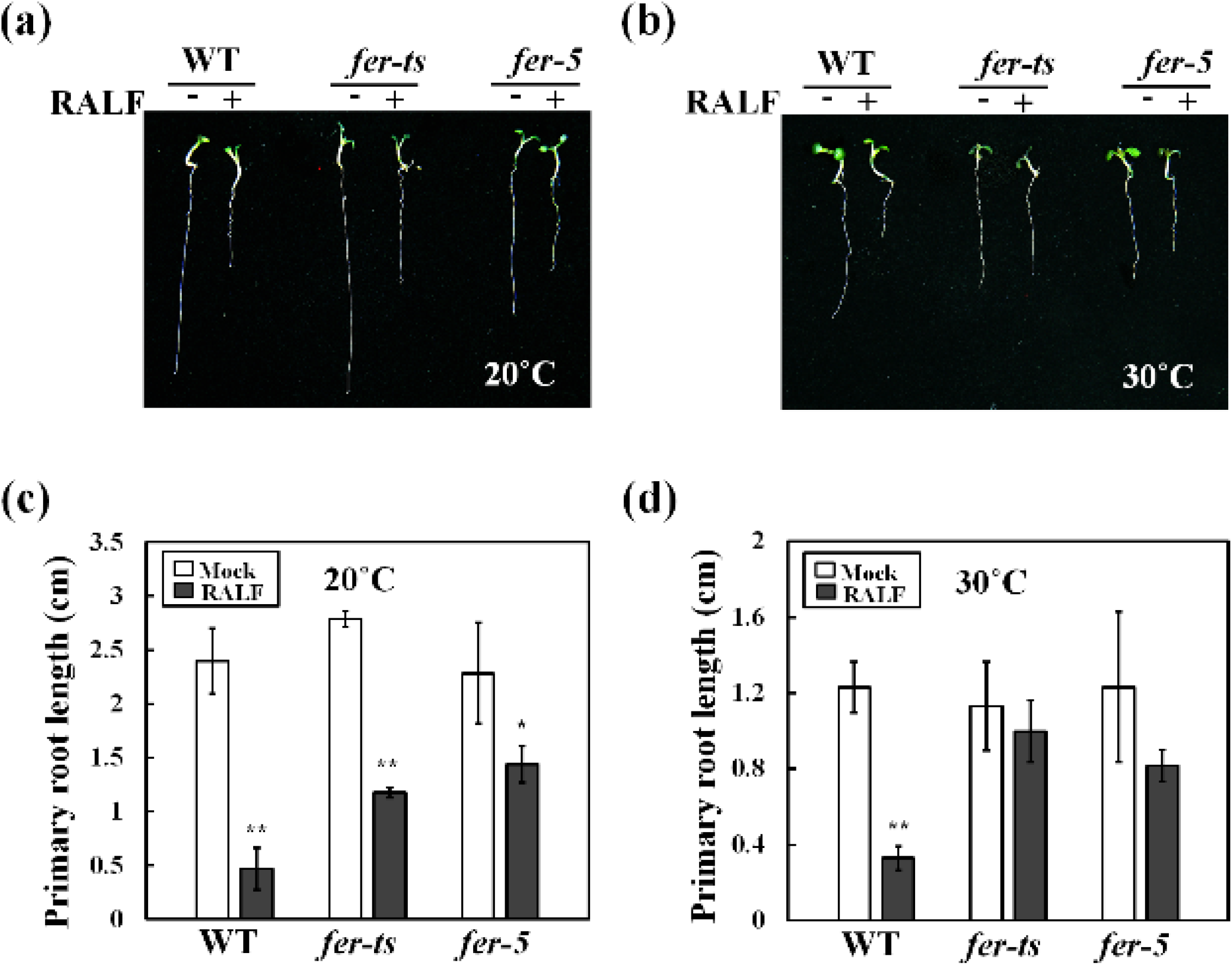
*fer-ts* mutants are partially insensitive to RALF1 peptide mediated root growth inhibition at elevated temperatures. Wild-type (WT), *fer-ts*, or *fer-5* plants were germinated and grown for 3 days in ½ MS liquid media at 20°C, and then transferred to ½ MS liquid media containing 1uM RALF1 peptide (RALF+) or a mock buffer control (RALF-) and grown an additional 3 days at 20°C (a) or 30°C (b). Images of representative seedlings were collected using an Olympus SZX12 stereoscopic microscope. Quantification of primary root lengths (n = 10 seedlings) in the presence or absence of RALF1 peptide treatment in normal, 20°C (c) and elevated, 30°C (d) conditions. Primary root lengths were determined using Image J. Error bars represent SD. *p<0.05, **p<0.01 by Student’s *t*-test.

### ROS accumulation of *fer-ts* was greatly reduced at 30 °C and *fer-ts* phenotype was not rescued by various hormone treatments

In previous reports, ROS accumulation is highly reduced in *fer-4* and *fer-5* mutants especially in root hair tips and primary roots (Duan et al., 2010). In order to investigate the ROS accumulation, WT and *fer* mutants were treated with H_2_DCF-DA to monitor ROS levels. In WT plants, ROS accumulation was observed in primary roots and root hairs, and in root hairs these levels increased slightly upon NAA treatment at both 20°C and 30°C (Figure 11). While *fer-ts* plants showed similar ROS accumulation patterns as those observed in wild-type plants at 20°C, at 30°C ROS accumulation was dramatically reduced both in the absence and presence of NAA (Figure 11a). However, while ROS accumulation in the *fer-ts* mutant was strictly temperature dependent (compare Figures 11b and 11c), these reduced ROS levels were similar to those observed at both temperatures for the constitutive *fer-4* and *fer-5* mutants (Figure 11a; compare Figures 11b and 11c)

**Figure 11.**
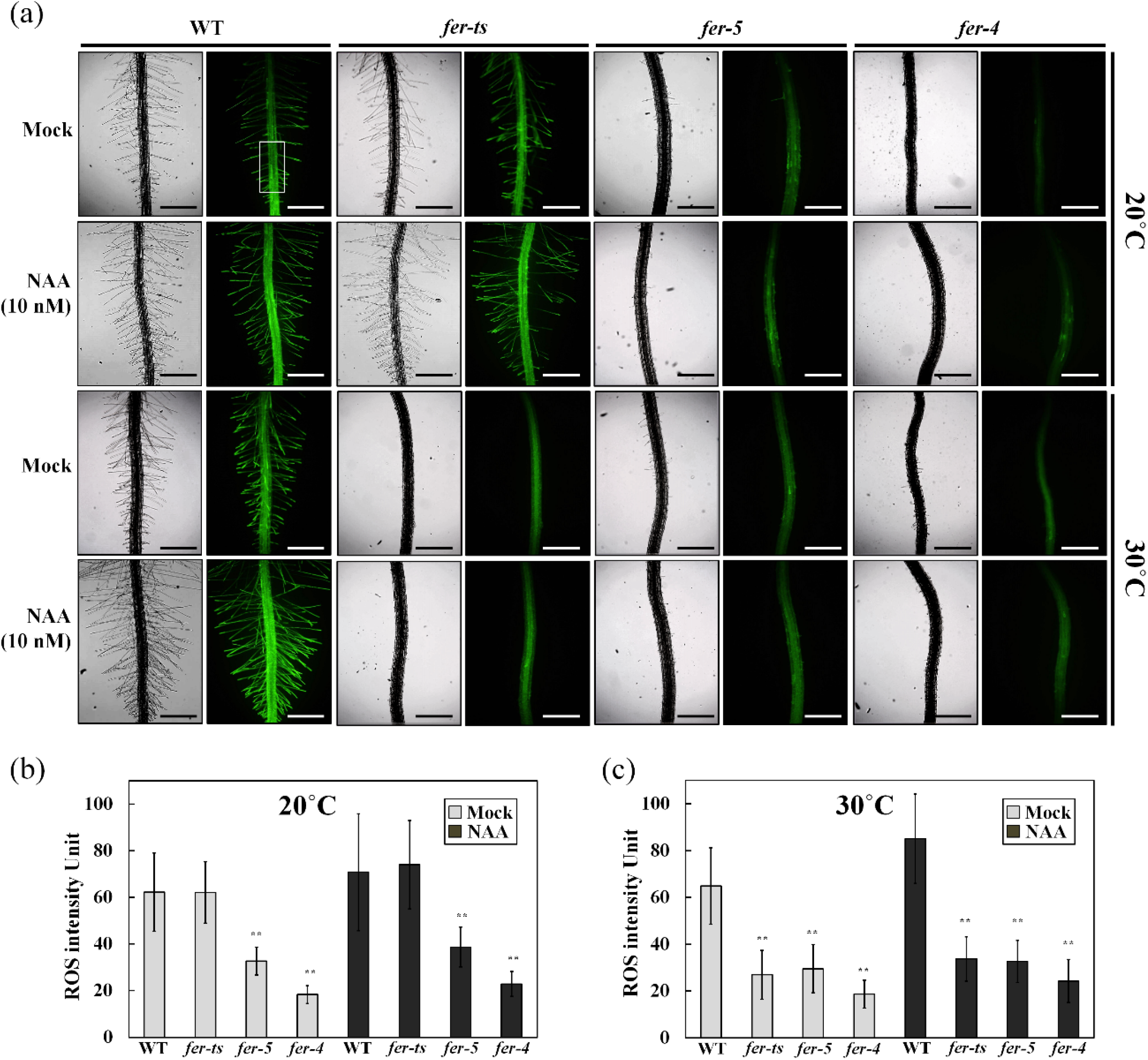
Detection of ROS in WT, *fer-ts* and *fer-5* primary roots and root hairs. (a) ROS accumulation in normal and elevated temperature conditions with or without auxin treatments. Wild-type (WT), *fer-ts*, *fer-4*, or *fer-5* seedlings were grown vertically on ¼ MS media plates for seven days at normal (20°C) or elevated (30°C) temperatures in the presence or absence of (10 nM NAA). Plates were bathed with five ml of 50 uM in H_2_DCF-DA suspended in ¼ MS liquid media for 5 min, followed by two gentle washes with 10 ml of ¼ MS. Fluorescence images were collected with a Zeiss Axio Imager Z1 fluorescence microscope with 2.5x objective and green (GFP) filter set. The WT ROS image was acquired by auto-exposure, all other images were acquired using the WT exposure conditions. Scale bars = 500 μm. (b) The rectangle in (a) indicates a representative region of interest (ROI) where average ROS intensity was quantified for the samples. Intensities of ROS were quantified by image J program. Error bars represent SD. **p<0.01 by Student’s *t*-test.

In order to investigate how broadly the temperature-sensitive *fer-ts* mutant phenocopied *fer-4* and *fer-5* mutants, these three mutants were treated with several different concentrations of hormones, and then root hair lengths and densities were measured either at 20°C or 30°C. Root hair lengths and densities displayed the similar patterns for both wild-type and *fer-ts* at 20°C, while both *fer-4* and *fer-5* root hairs were consistently shorter and less dense (Supplemental Figure S5a and S5b; upper panels). However at 30°C, *fer-ts* root hair lengths and densities largely resembled the *fer-4* and *fer-5* phenotypes (Supplemental Figure S5a and S5b; lower panels). Primary root length, total lateral root number, fresh weight, and total leaf numbers of *fer-ts* mutants also displayed a similar temperature-dependent trend; resembling wild-type plants with various hormone treatments at 20°C, but resembling *fer-4* and *fer-5* mutants at 30°C (See supplemental Figure S6).

## DISCUSSION

In eukaryotes, receptor like kinases (RLKs) have been implicated to play an important role in many crucial eukaryotic cellular processes, such as cell cycle progression, cell signaling, embryogenesis, abiotic and biotic stress responses (Shiu and Bleecker, 2001; Morillo and Tax, 2006; Lehti-Shiu et al., 2009). In this study, we isolated and identified a temperature-sensitive root hair elongation mutant, which we have determined is a new mutant *FER* allele that we have called *fer-ts*. The *fer-ts* mutant displays normal overall growth characteristics at normal temperature (20°C), but root hair initiation and elongation are specifically and rapidly inhibited within one minute upon transfer of these plants to elevated temperature (30°C). We have shown that the *fer-ts* mutant is the result of a substitution mutation in which a highly conserved glycine residue in the FER extracellular domain is changed to serine (G41S). FERONIA is a member of the CrRLK 1 L subfamily of receptor-like kinases (RLKs) in *Arabidopsis* and the mutated glycine residue (G41S) is highly conserved in multiple members of the CrRLK1L family of receptor proteins as well as in animal malectin proteins.

While both FER(WT)-EYFP and FER(G41S)-EYFP fusion proteins displayed a plasma membrane localization at both normal and elevated temperatures, FER(G41S)-EYFP displayed significantly increased protein turnover 30°C (Figure 6). However, this increased protein turnover did not appear to result in loss of accumulation of FER(G41S)-EYFP in plasma membranes. This is likely due to continued protein synthesis and at least some secretion of these proteins at the elevated temperature. These results, along with the rapid cessation of root hair elongation (<1 min) in elevated temperatures would be consistent with rapid inactivation of FER signaling activity due to protein inactivation rather than simply depletion of FER activity from the plasma membrane due to increased turnover.

*FER* has been implicated in a variety of plant processes, including roles in root hair tip growth as well as crucial plant processes, such as pollen tube reception, hypocotyl elongation, regulation of ABA signaling and controlling seed size (Escobar-Restrepo et al., 2007; Deslauriers and Larsen, 2010; Duan et al., 2010; Yu et al., 2012; Yu et al., 2014). In many of these processes, FER signaling appears to regulate ROS production. In constitutive *fer* mutants, ROS levels are reduced, and FER overexpression results in increased ROS levels. The observation that the *fer-ts* mutant also displays reduced ROS levels only at elevated temperatures suggests that this mutation affects FER signaling in a similar fashion as other *fer* mutants, perhaps providing a powerful tool for elucidation of downstream signaling events associated with FER function, and indicating that at least one important downstream effect of *FER* signal transduction is regulation of ROS production. This was elegantly explained by the discovery that FER recruits ROPGEFs, which in turn activate ROP GTPases, leading to the stimulation of RHD2 NADPH oxidase dependent ROS production (Duan et al., 2010). Therefore, FER mediated regulation of ROS production is likely important and tightly controlled for many cellular functions.

Based on sequence comparison, the extracellular domains of members of the CrRLK 1 L subfamily of plant RLK proteins were predicted to share some structural similarity to the mammalian malectin protein (Schallus et al., 2008). Malectin was first identified and characterized in *X. laevis* as carbohydrate binding protein of the endoplasmic reticulum where it plays an important role in the early steps of protein N-glycosylation for biogenesis of glycoproteins (Schallus et al., 2008). Based on NMR structure analysis, there are five key residues in the malectin domain (Y67, Y89, Y116, F117, D186) that are located in pocket-shaped structure and these aromatic residues and the aspartate mediate interactions with the glucose residues of maltose and nigerose disaccharide ligands (Schallus et al., 2008). In plants, malectin-like domains are mainly found in CrRLK1L subfamily with a low overall sequence identity with animal malectins (Shiu and Bleecker, 2003). In FER, two malectin-like domains, ML1 and ML2, are found as a tandem-repeat in the extracellular domain. Interestingly, several key residues found in the ligand-binding pocket of the animal malectin structure are maintained in the malectin-like domains of FER and other plant CrRLK1L family members (Schallus et al., 2008). However, the discovery that members of a family of small secreted peptides, RALFs, rather than cell wall polysaccharides or oligosaccharides, serve as important ligands for FER and other CrRLK1L family receptors (Haruta et al., 2014; Ge et al., 2017; Stegmann et al., 2017; Gonneau et al., 2018) might indicate that these extracellular domains may interact with ligands in a manner distinct from their animal counterparts. Indeed the recent structural characterization of ANX1/2 extracellular domains (Du et al., 2018) and the FER extracellular domain in complex with RALF23 and the FER coreceptor, LLG2 (Xiao et al., 2019) has shown that the RALF23 binding domain and interaction with LLG2 occurs primarily with the ML2 domain, and that conserved tyrosine and phenylalanine residues in CrRLK1L malectin folds in these structures appear to be buried within the ML1 fold, and therefore likely unavailable to interact with cell wall carbohydrates in a manner similar to animal malectins.

On the other hand, analysis of the animal and plant malectin domains, reveals an additional invariant glycine residue, that is present in all animal and plant malectin sequences, and which is also found in close proximity to pocket-shape ligand-binding cleft determined in the structure of the animal ML1 domain. This invariant glycine is replaced with a serine (G41S) in the *fer-ts* mutation described in this paper. The highly conserved nature of this glycine residue, and the rapid elimination of FER signaling at elevated temperatures, suggests a critical role for the FER ML1 domain in ligand binding or transduction of a ligand-binding signal in members of the CrRLK1L family of receptor-like kinases. Indeed, mutation of an analogous glycine residue to aspartic acid (G37D) in the extracellular domain of THESEUS in the *the1-1* mutant also results in a loss of function mutation in this RLK (Hematy et al., 2007). The *the1-1* mutation also results in its insensitivity to its specific RALF ligand, RALF34 (Gonneau et al., 2018). However, RALF34 was still found to bind the extracellular domain of *the1-1* containing the G37D mutation in vitro, suggesting that mutation of this glycine residue did not directly impact RALF34 binding (Gonneau et al., 2018). Similarly, the response of *fer-ts* mutant to treatment with RALF1 peptide was dramatically reduced under elevated temperature conditions (Figure 7). Precisely how the G41S *fer-ts* mutation, which is not found within the RALF23 peptide binding surface in the FER ML2 domain, would directly block RALF peptide perception and signaling is unclear. One potential explanation may involve the recent discovery of links between *FER* signaling and pectin dynamics during salt stress (Chen et al., 2016; Feng et al., 2018) and fertilization events (Duan et al., 2020). During salt stress, *FER* appears to sense cell wall softening and both FER ML1 and FER ML2 domains were shown to directly interact with pectin *in vitro* (Feng et al., 2018). More recently, *FER* function was shown to be required in order to maintain de-esterified pectin levels in the filiform apparatus during pollination and fertilization events (Duan et al., 2020), Whether the G41S mutation in *fer-ts*, or other analogous mutations of this invariant glycine residue in other CrRLK1L receptors affect the ability of these receptors to interact with or regulate pectin dynamics in plant cell walls is an intriguing possibility that warrants future investigation.

## EXPERIMENTAL PROCEDURES

### Plant materials and growth conditions

*Arabidopsis thaliana* ecotype Columbia (Col-0), the *fer-ts* mutant was isolated from an EMS mutagenized population of wild-type (Col-0) stably transformed with a single copy of EYFP-RabAb4 driven by a 35S promoter (Preuss et al., 2004; Weigel and Glazebrook, 2006), and two FERONIA T-DNA insertional mutants (designated as *fer-ts* and two T-DNA insertion mutants, *GABI-GK106A06* (designated as *fer-4*) and *SALK_029056c* (designated as *fer-5*) (Duan et al., 2010) were used in this study. Seeds were sterilized by soaking in 1% bleach solution for 10 min; after washing five times with sterilized water, they were sown onto agar plates for germination. Five-to seven-day-old *A. thaliana* seedlings used in the root-hair growth assays were grown vertically on plates containing 0.25x Murashige and Skoog (MS; Sigma-Aldrich) medium at pH 5.7 supplemented with 0.6% (w/v) phytagel at 20°C under long day conditions (16-h days with 150 μE·m^-2^·s^-1^ (E, Einstein; 1 E = 1 mol of photons). For harvesting seeds, seedling plants were transferred to soil and grown to maturity at 20°C under long day conditions.

### Quantification of root hair elongation in *fer-ts* under normal and elevated temperatures

To characterize root hair growth defective phenotypes in the *fer-ts* mutant, bright-field microscopy was carried out using a Nikon Eclipse E600 wide-field microscope with a 10x Plan Apo (0.45 NA) lens as previously described in (Preuss et al., 2004). The *fer-ts* mutants were germinated and grown vertically on plates containing 0.25x MS medium at pH 5.7 supplemented with 0.6% (w/v) phytagel at 20°C for 7 days and then transferred to 30°C and grown for 6 hours before returning back to 20°C growth conditions for an additional 24 hours, when images of roots and root hairs were then collected. Time-lapse video microscopic analysis was carried out under normal and elevated temperature conditions in wild-type and *fer-ts* mutants as described previously (Preuss et al., 2004). Images of growing root hairs were collected from seedlings by time-lapse video microscopy either using a Nikon Eclipse E600 wide-field microscope with a 10x Plan Apo (0.45 NA) lens (for bright-field images), or using a Leica confocal laser-scanning microscope SP8 with a 63x oil lens (1.4 NA). YFP fluorescence was excited at 514 nm and visualized from 519 nm to 650 nm (Yang et al., 2020). The temperatures of MS medium was controlled by an inline single-channel automatic temperature controller (Werner Instruments, Hamden, CT, model:TC-324B) controlled by a dry air thermostat inserted into the growth chamber and situated approximately 2 mm from the ROI. Temperatures were actively recorded using an Infrared Thermometer (Kintrex Inc., Vienna, VA, model:IRT0424). Raw image sequences were cropped with Adobe Photoshop and imported into Fiji-ImageJ (Schindelin et al., 2012) to generate time projections using the Stacks function. Quantification of root-hair lengths, growth rates, and densities were quantified by using calibrate and measure functions.

### Map-based cloning and full genome sequencing of *fer-ts*

Self-fertilized, backcrossed *fer-ts* (ecotype; Columbia) mutants were crossed with Landsberg wild-type plants to generate a mapping population. F1 crossed plants were checked for heterozygosity with the SSLP marker “nga8” that is polymorphic between Col-0 and L*er* (Bell and Ecker, 1994). Homozygous *fer-ts* plants were selected from the segregating F_2_ population by germination on MS media plates at 20°C, and subsequent analysis of root hair tip growth defective phenotypes in elevated growth temperatures (30°C). Homozygous plants displaying *fer-ts* phenotypes were grown to maturity at 20°C and seed were collected. Genomic DNA was isolated using Qiagen Plant DNA mini kits, and SSLP markers were used for rough mapping the *fer-ts* mutant lesion, which was initially located on chromosome 3 between the SSLP markers *NIT 1.2* and *NGA6*. Low-resolution mapping narrowed the location of the *fer-ts* mutant locus to an approximately 2 Mb region of chromosome 3, and full genome sequencing was performed to further determine the *fer-ts* mutation within this region. Libraries were generated for both the *fer-ts* and wild-type (ecotype Columbia) extracted DNA using Illumina TruSeq DNA kits and barcoded for multiplexing by the University of Michigan DNA Sequencing Core. Samples were sequenced on an Illumina MiSeq platform with paired-end 150 bp cycles. Sequence reads were checked for quality using FastQC then aligned to the TAIR9 genome using Bowtie2. Potential SNPs were identified using Freebayes. Additional analysis of sequence variants within the low-resolution mapped 2 Mb region of chromosome 3 to eliminate SNPs common to our re-sequenced Col-0 population and the *fer-ts* allele were sorted for context and predicted effect using a custom PERL script.

### Fluorescence microscopic analysis

For pFER-FER(WT)-EYFP and pFER-FER(G41S)-EYFP transgenic plants, full-length of *FERONIA* including approximately 2 kb promoter was prepared by PCR reaction and sub cloned into the pCAMBIA-EYFP-C1 expression vector (Preuss et al., 2004). Wild-type and mutant FERONIA sequences were amplified from genomic DNA isolated from wild-type and *fer-ts* mutant plants using PCR. To produce pFER-FER(WT)-EYFP and pFER-FER(G41S)-EYFP transgenic plants, these constructs were introduced into *A. tumefaciens* strain GV3101, and *Arabidopsis* plants were transformed with *A. tumefaciens* using the ‘floral-dip’ method (Clough and Bent, 1998). Transgenic plants for each construct were selected by germination on 25 mg/L of hygromycin-containing medium (Duchefa, Haarlem, The Netherlands) under long day conditions (16 h light/8 h dark cycle) at 20°C. Three or more independent, homozygous transgenic lines were isolated for each of the pFER-FER(WT)-EYFP and pFER-FER(G41S)-EYFP constructs in the *fer-ts*, *fer-4*, and *fer-5* mutant backgrounds, respectively, and a single representative homozygous line was used in these studies. Confocal images were generated using a laser confocal microscope (Zeiss Observer.A1) connected to a CSU10 confocal scanner unit (Yokogawa, Japan) and a 10x Plan-Neofluar (0.3 NA lens), 40x Plan-Apochromat (1.3 NA lens) or 100x Plan-Apochromat (1.46 NA lens) oil objective with 491 nm laser excitation and a 535 nm emission filter for EGFP and EYFP fluorescence. To visualize cell walls seedlings were incubated for five minutes in Propidium Iodide (10 μg/ml) and then washed with 0.25xMS media, cell membranes were stained with FM 4-64 (5 μM) for five minutes before washing with 0.25xMS media. Images were collected with a Hamamatsu C9100-50 camera operated using the Volocity software version 5 (the electron-multiplying (EM)-CCD detector gain settings were 123, 116 and 190 for images collected with 10x, 40x and 100x objectives, respectively).

### RT-PCR analysis

For detection of *FERONIA* expression in wild-type, *fer-4, fer-5* and *fer-ts* plants, plants frozen immediately in liquid nitrogen. Two microgram aliquots of total RNA extracted from the wild-type or mutant seedlings were used for reverse transcription primed by oligo(dT). Superscript III (Invitrogen, USA) was used for the reverse transcription reaction according to the manufacturer’s instructions. One microliter aliquot of the reaction mixture was used for subsequent PCR analysis. *Actin* was used as a quantifying control.

### Detection of ROS in roots

ROS detection by using H_2_DCF-DA in root hair and primary root was performed following the protocol described previously (Duan et al., 2014). Briefly, Arabidopsis seedlings were germinated grown vertically on 0.25xMS media plates for seven days at normal (20°C) or elevated (30°C) temperatures. Plates were bathed with five ml of 50 uM in H_2_DCF-DA (Sigma-Aldrich) suspended in 0.25xMS liquid media for 5 min, followed by two gentle washes with 10 ml of 0.25xMS. Fluorescence images were collected with a Zeiss Axio Imager Z1 fluorescence microscope with 2.5x objective and green (GFP) filter set.

### Effect of RALF1 peptide on root hair and root growth inhibition

Synthetic Arabidopsis RALF1 polypeptide was synthesized by using 9-fluorenylmethyl chloroformate solid-phase chemistry with a peptide synthesizer from Thermo Scientific company and confirmed by MALD-TOF analysis (Applied Biosystems Voyager System 2098, USA). After synthesis, 5 mg of reduced synthetic polypeptide was oxidized by dissolving in 25 ml of degassed 0.1 M ammonium bicarbonate and incubating for 2 days in an opened flask under 4°C, then lyophilized. Lyophilized RALF1 powder was re-suspended in 10 ml of PBS buffer followed by two buffer exchange steps using Amicon Ultra centrifugal filter (Ultracel-3K, 3000g for 45 min each) to remove any residual ammonium bicarbonate. Seedling germination was performed in 0.5xMS liquid medium at 20°C for 3 days in long-day conditions (16-h days with 150 μE·m^-2^·s^-1^ (E, Einstein; 1 E = 1 mol of photons) light intensity. After 3 days, germinated Arabidopsis seeds were transferred to 6 well Falcon tissue culture plate with 3 ml of 1/2xMS liquid media containing 1 μM RALF1 or an equal volume of PBS and agitated on a shaker at 100 rpm (Model VS2010, Vision Scientific CO.,LTD) for an additional 3 days at 20°C or 30°C. All solutions were filter-sterilized (0.2 μm pores, Minisart 16534), and the seedlings were photographed 3 days after being transferred to the media. Quantification of primary root length was measured using ImageJ software. Time-lapse video microscopic analysis was carried out under permissive temperature conditions in wild-type. 0.25x MS liquid media with/without 1 μM RALF1 peptide were pumped to the chamber while imaging. All solutions were filter-sterilized (0.2 μm pores, Minisart 16534).

### Accession numbers

Sequences of the genes in this paper may be found in the GeneBank/EMBL database library under the following accession numbers: At3g51550 (FER), At4g39990 (RabA4b), At3g18780 (Actin2), At3g04690 (ANX1), At5g28680 (ANX2), At3g46290 (HERK1), At1g30570 (HERK2), At5g54380 (THE1), NP_001085212.1 (*X. laevis* Malectin), NP_055545.1 (*H. sapiens* Malectin).

## ACKNOWLEDGEMENTS

We thank to Dr. Hen-Ming Wu and Dr. Alice Cheung (University of Massachusetts) for providing *fer-4* and *fer-5* mutants, and Jiyuan Yang for assistance in assembling and editing the manuscript. This research was supported by the U.S. Department of Energy Office of Science, Office of Basic Energy Sciences, Physical Biosciences program (DE-FG02-07ER15887; S.P., F.G., J.C., and E.N,), the National Science Foundation under Grant No. 1817697 (A.A., H.M, and E.N.), and the BK21plus program of the Ministry of Education, Science and Technology in Korea (D.K., and J-D.B.). This work used the Extreme Science and Engineering Discovery Environment (XSEDE; (John W Towns, 2014), which is supported by National Science Foundation grant number ACI-1548562.

## SUPPORTING INFORMATION

Additional supporting Information may be found in the online version of this article.

**Figure S1.**
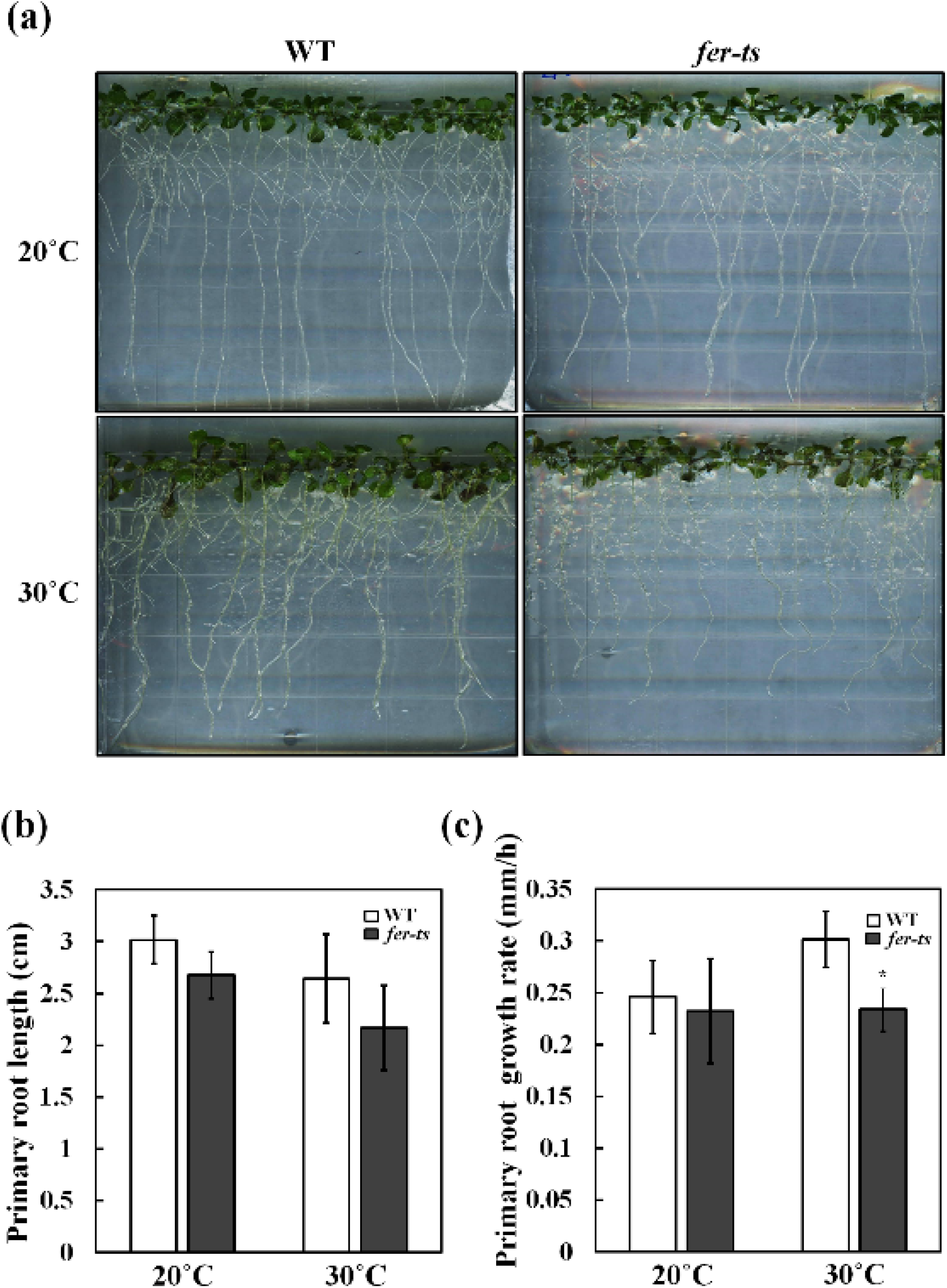
Primary root growth of *fer-ts* mutants under normal and elevated temperature conditions. (a) Photographs of wild-type and *ltl2* mutant under permissive and non-permissive temperature conditions. (b) Measurements of primary root growth length under normal (20°C) and elevated (30°C) temperature conditions. (c) Primary root growth rate under 20°C and 30°C conditions. (a)-(c) Seven day-old wild-type (WT; n=25) and *fer-ts* (n=25) seedlings were transferred to the 30°C for two weeks and primary roots were scanned by EPSON 4990 photo scanner every second day. The primary roots length were determined using Image J, and root growth rates were calculated by dividing growth by 48 hours (mm/h). Error bars represent SD. *p<0.05 by Student’s *t*-test.

**Figure S2.**
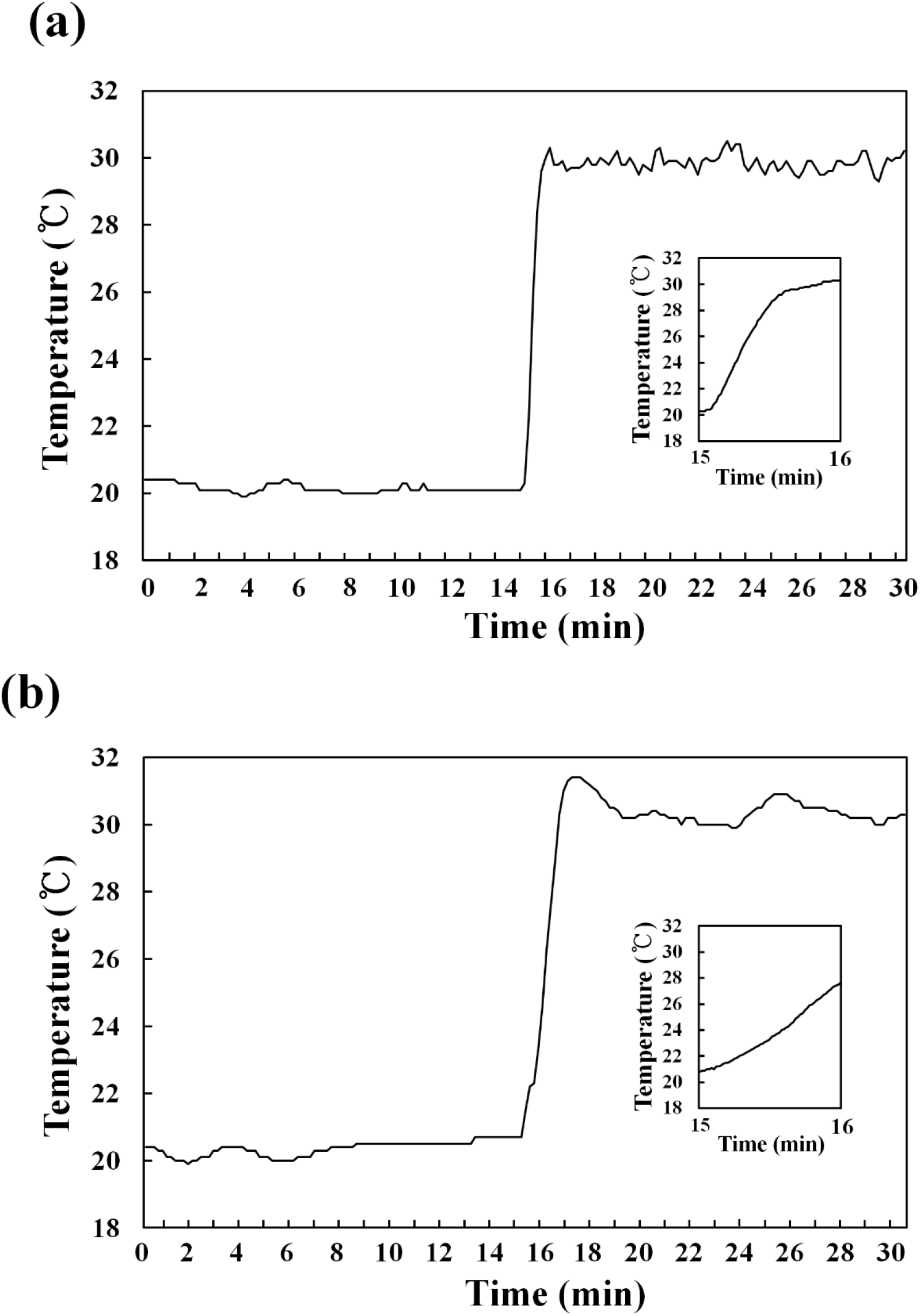
Root hair growth chamber temperature measurement during transition from normal (20°C) to elevated (30°C) temperature conditions.

**Figure S3.**
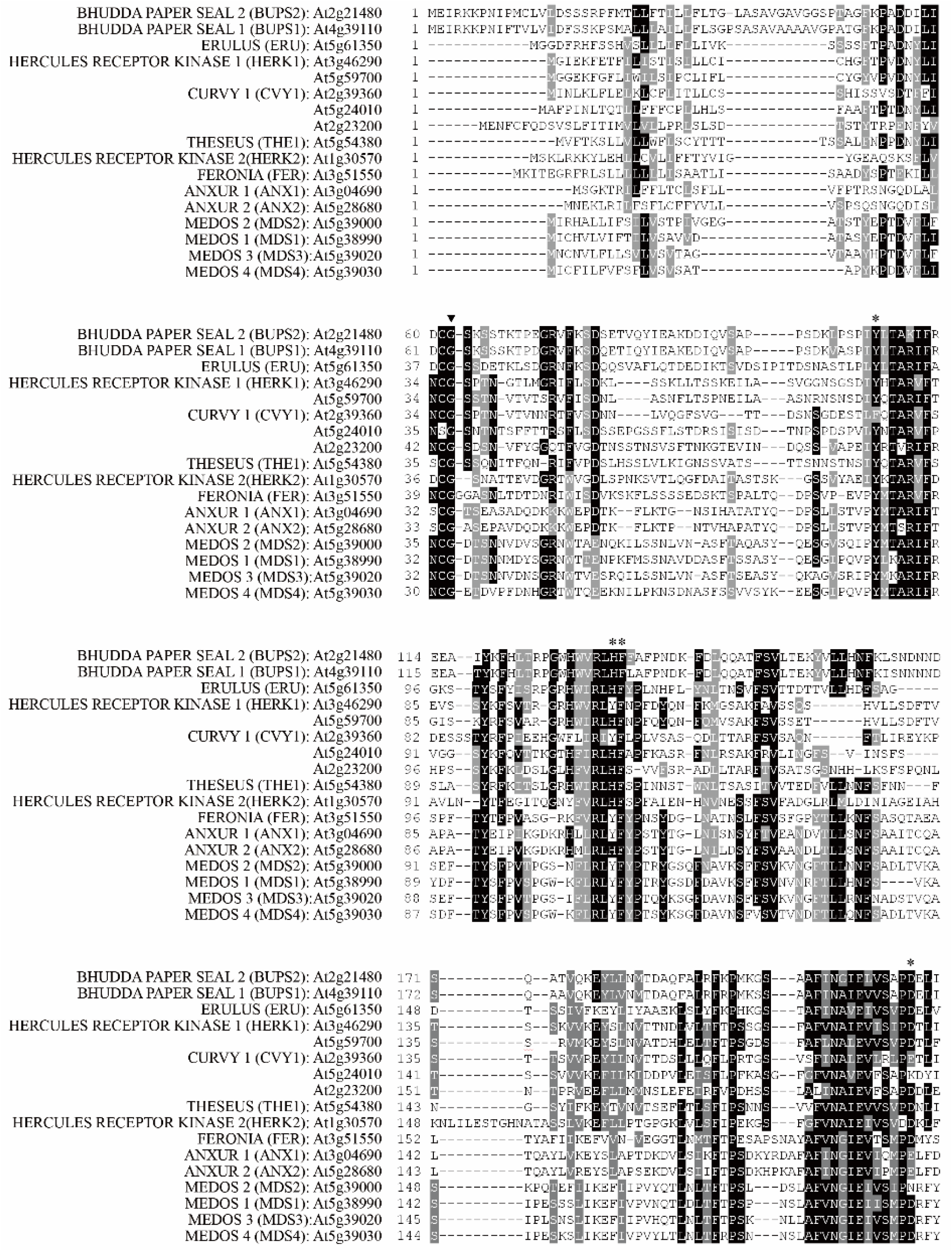
Sequence alignment of *Arabidopsis* CrRLK1L subfamily receptor kinases. Multiple-sequence alignment analysis was performed with the Clustal Omega program (http://www.ebi.ac.uk/Tools/msa/clustalo/) and displayed using BOXSHADE (www.ch.embnet.org/software/BOX_form.html). Sequence similarity analysis of CrRLK subfamily members in Arabidopsis. *At3g51550* encodes FERONIA protein. The highly conserved FERONIA G41 and additional putative key residues are marked by black arrowheads and asterisks, respectively. Black boxes indicate conserved residues; shaded boxes indicate residues with conserved substitutions.

**Figure S4.**
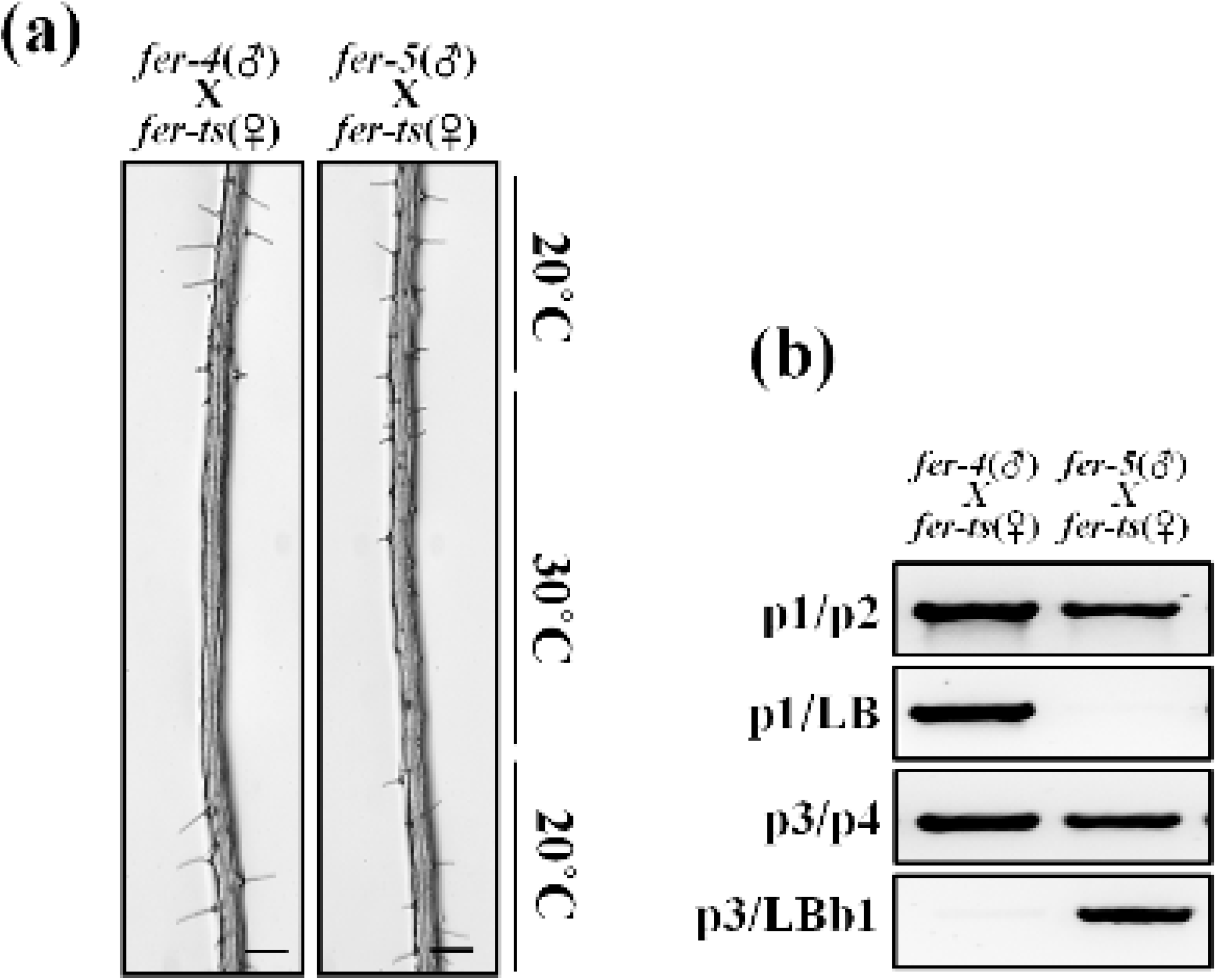
Temperature-sensitive root hair growth in *fer-ts/fer-4* and *fer-ts/fer-5* F1 plants. (a) Wild-type (WT), *fer-4, fer-5*, and F1 progeny from crosses (maternal = *fer-ts*, paternal = *fer-4* or *fer-5*) of *fer-ts/fer-4* and *fer-ts/fer-5* were grown vertically for seven days at 20°C, transferred to 30°C for 6 h, and then grown for an additional 24 h at 20°C. Both *fer-ts/fer-4* and *fer-ts/fer-5* progeny clearly demonstrated a ts-dependent root hair phenotype. Scale bars = 200 μm. (b) Genotyping of crossed F1 plants. Genomic DNA was extracted from F1 generation plants and subjected PCR to confirm presence of the *fer-4* and *fer-5* genotypes.

**Figure S5.**
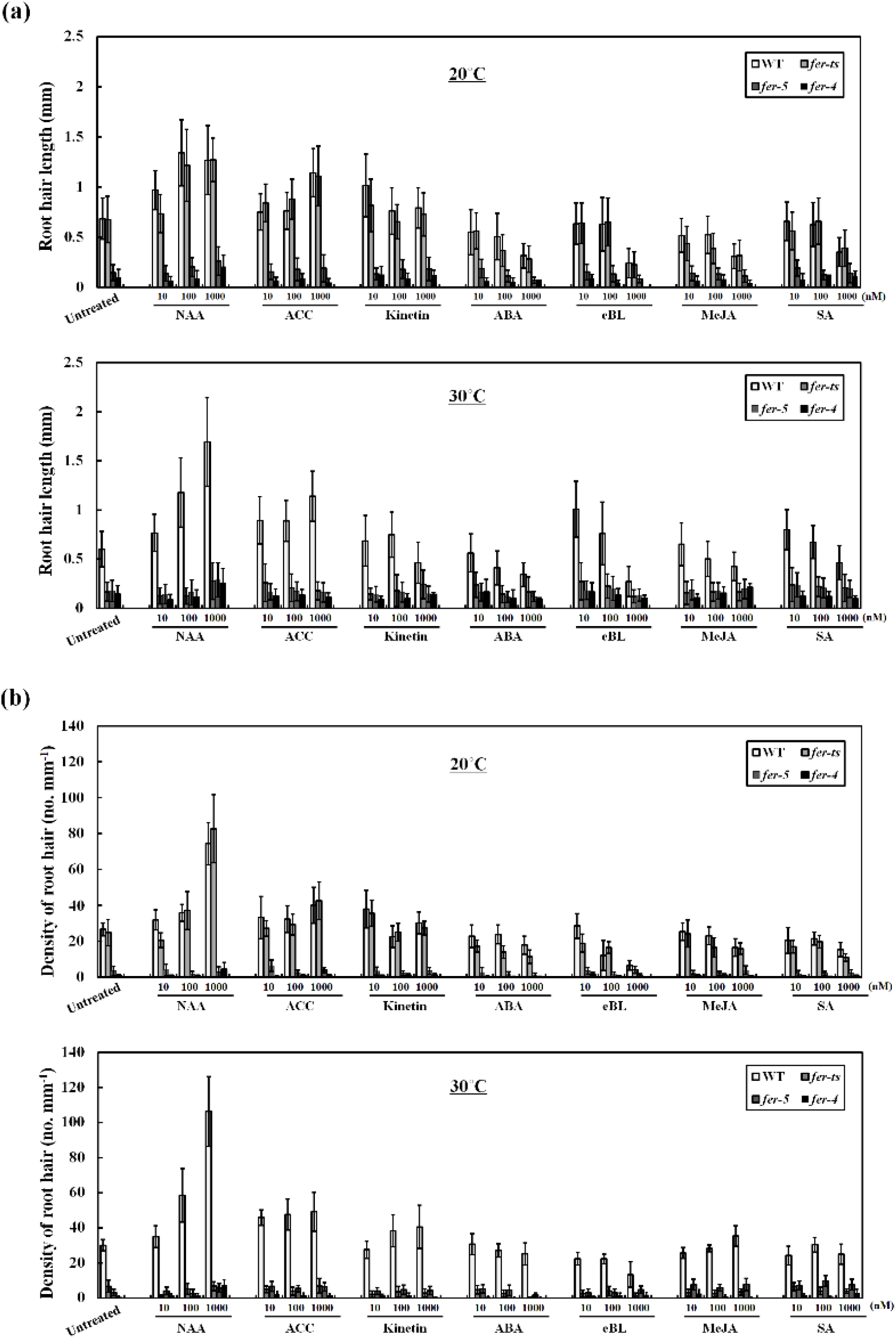
In elevated temperatures, *fer-ts* responds to hormone treatments similarly to *fer-4* and *fer-5* mutants. (a and b) Quantification of root hair length and density in normal (upper panels) and elevated temperature (lower panels) in 10, 100, and 1000 uM concentrations for each of the various hormone treatments (NAA; Auxin, ACC; ethylene, Kinetin; Cytokinin, ABA; Abscisic acid, eBL; epi-brassinosteroid, MeJA; methyl jasmonic acid, SA; Salicylic acid). Three-day-old seedlings were transferred into ½ MS liquid media containing three different concentrations of the various hormones. Then, transferred plants were incubated at 20°C and 30°C for seven days before quantification. The root hairs were photographed by Olympus AX-70 microscope and root hair length and densities were determined by image J. Error bars represent SD.

**Figure S6.**
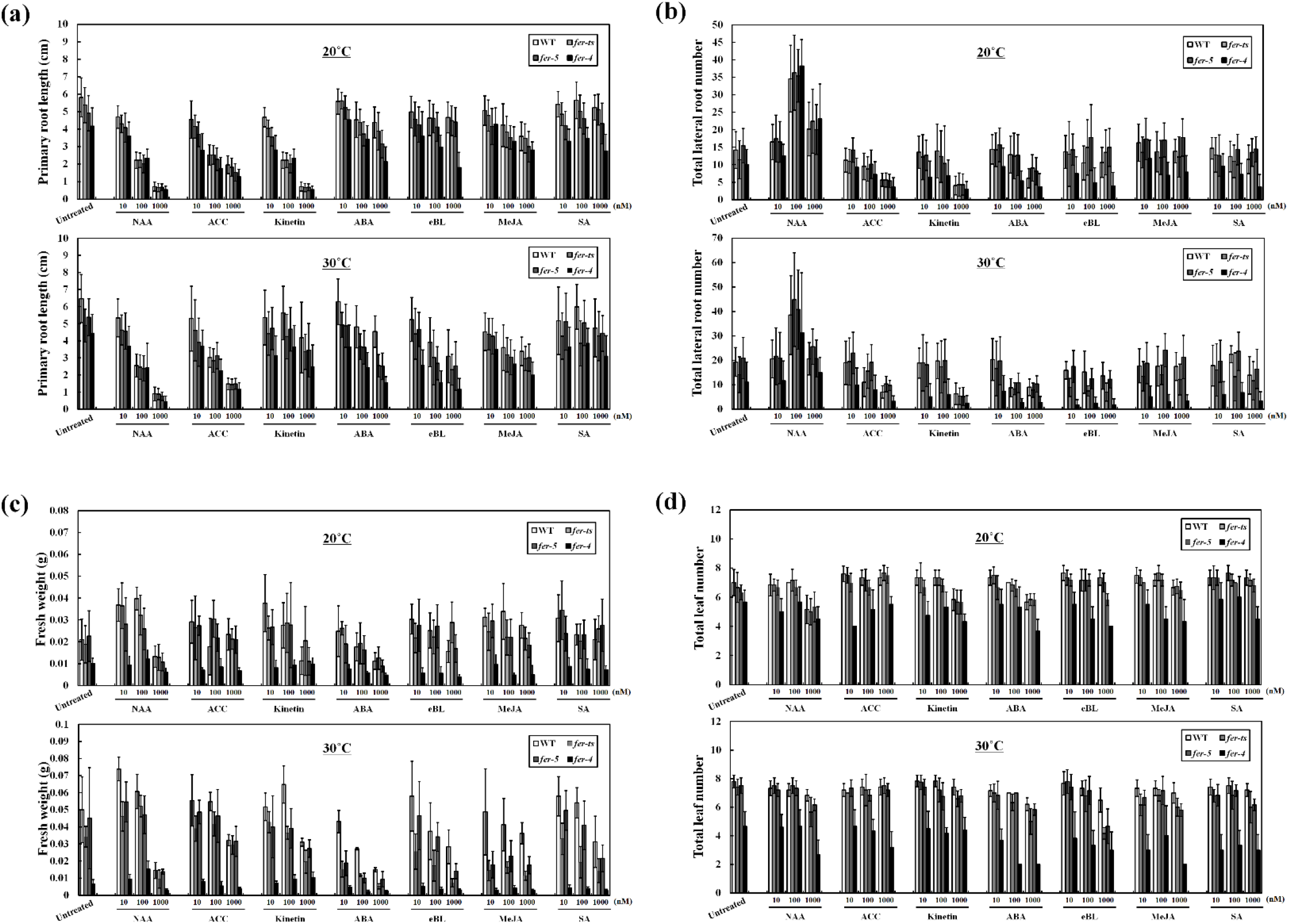
Hormonal effects on primary root length, total lateral root number, fresh weight and total leaf number in normal and elevated temperature conditions for *fer-ts, fer-4*, and *fer-5* plants. Primary root length (a), lateral root number (b), fresh weight (c), and leaf number (d) were measured in normal (upper panels) and elevated (lower panels) temperatures in 10, 100, and 1000 uM concentrations for each of the various hormone treatments (NAA; Auxin, ACC; ethylene, Kinetin; Cytokinin, ABA; Abscisic acid, eBL; epi-brassinosteroid, MeJA; methyl jasmonic acid, SA; Salicylic acid). Three-day-old seedlings (n=20 each for wild-type, *fer-ts*, *fer-4*, and *fer-5*) were transferred into ½ MS liquid media containing three different concentrations of the various hormones. Then, transferred plants were incubated at 20°C and 30°C for seven days before quantification. Error bars represent SD.

**Movie S1.** Time-lapse imaging of growing root hair of WT and *fer-ts* mutants during permissive and non-permissive temperatures. The root hair growth of WT and *fer-ts* plants (seven day-old seedlings) mounted in a temperature-controlled perfusion chamber were imaged with a Nikon ECLIPSE E600 microscopy with a 10x Plan Apo (0.45 NA) lens. Perfused plant growth chamber temperatures were controlled by a combination of a dry air thermostat linked to a single channel Automatic Temperature Controller TC-324B, and real-time measurement using an Infrared Thermometer IRT0421. Images of WT and *fer-ts* mutant root hairs were collected every five seconds and cropped by Photoshop Lightroom version 5.3. Video clips were constructed by VEGAS 6.0 program from Sony (30 frames/s).

**Movie S2.** Dynamics of YFP fused FERONIA protein localization in growing root hair. The growing root hair of YFP fused FERONIA protein (seven day-old seedlings) using perfusion chambers was observed by Zeiss Observer.A1 laser confocal microscopy connected to a CSU10 confocal scanner unit (Yokogawa, Japan) and a 63x Plan-Apochromat (1.40 NA) oil objective with 491 nm laser excitation and a 535 nm emission filter for EYFP fluorescence. Images were collected with a Hamamatsu C9100-50 camera and operated using the Volocity software version 5. Time-lapse fluorescent images were taken every 5 s for 20 min and automatically constructed by Volocity software version 5 (30 frames/s).

**Movie S3.** FER(WT)-EYFP and FER(G41S)-EYFP protein localization in root hairs at normal (20°C) and elevated (30°C) temperature conditions. Subcellular localization of pFER-FER(WT)-EYFP and pFER-FER(G41S)-EYFP fusion proteins in root hair cells of seven-day-old seedlings were collected ever 5 s for 350 s at 20°C at which point the perfusion chamber was transferred to 30°C for an additional 300 s. Perfused plant growth chamber temperatures were controlled by a combination of a dry air thermostat linked to a single channel Automatic Temperature Controller TC-324B, and real-time measurement using an Infrared Thermometer IRT0421. Confocal images were collected YFP signal were acquired using a Leica confocal laser-scanning microscope SP8 with a 63x oil lens (1.4 NA). Movies were constructed using ImageJ (20 frames/s).

**Movie S4.** Effect of RALF1 peptide on root hair growth. Bright-field images of root hair cells of seven-day-old seedlings in a stably-transformed *fer-4* line expressing FER(WT)-EYFP were imaged using a Nikon Eclipse E600 wide-field microscope with a 10x Plan Apo (0.45 NA) lens. Time-lapse images were collected every 5 s for 20 min at 20°C in 0.25xMS media, at which point the perfusion buffer was switched to 0.25xMS media with 1 μM RALF1 peptide. Movies were constructed using ImageJ (20 frames/s).

**Table S1. List of primers used in this study.**

